# “A negative feedback loop mediated by the NR4A family of nuclear hormone receptors restrains expansion of B cells that receive signal one in the absence of signal two”

**DOI:** 10.1101/2020.03.31.017434

**Authors:** Corey Tan, Ryosuke Hiwa, James L. Mueller, Vivasvan Vykunta, Kenta Hibiya, Mark Noviski, John Huizar, Jeremy Brooks, Jose Garcia, Cheryl Heyn, Zhongmei Li, Alexander Marson, Julie Zikherman

**Author notes:** Address correspondence to: Dr. Julie Zikherman, 513 Parnassus Avenue, Room HSW1201E, Box 0795, San Francisco, CA 94143-0795. These two authors contributed equally to this manuscript (Corey Tan, Ryosuke Hiwa).

## Abstract

Ag stimulation (signal 1) triggers B cell activation and proliferation, and primes B cells to recruit, engage, and respond to T cell help (signal 2). However, failure to receive signal 2 within a defined window of time results in an abortive round of proliferation, followed by anergy or apoptosis. Although the molecular basis of T cell help has been extensively dissected, the mechanisms that restrain Ag-stimulated B cells, and enforce dependence upon co-stimulation, are incompletely understood. *Nr4a1-3* encode a small family of orphan nuclear receptors that are rapidly induced by B cell receptor (BCR) stimulation, yet little is known about their function in humoral immune responses. Here we use germline and conditional loss-of-function mouse models to show that *Nr4a1* and *Nr4a3* play partially redundant roles to restrain both the survival and proliferation of B cells that receive signal 1 in the absence of co-stimulatory signals, and do so in part by repressing expression of BATF and consequently c-MYC. Correspondingly, Ab responses to TI-2 immunogens are enhanced in the absence of *Nr4a1*, but are unaltered in response to immunogens that incorporate co-stimulatory signals. Unexpectedly, we also identify a role for the NR4A family in restraining B cell access to T cell help by repressing expression of the T cell chemokines CCL3/4, as well as CD86 and ICAM1, and show that this is relevant under conditions of competition for limiting T cell help. Our studies collectively reveal a novel negative feedback loop mediated by the NR4A family that increases B cell dependence upon T cell help and restrains strongly Ag-activated B cell clones from monopolizing limiting amounts of T cell help. We speculate that this imposes B cell tolerance and dampens immunodominance to facilitate preservation of clonal diversity during an immune response.

## INTRODUCTION

The requirement for two distinct types of stimuli to fully activate B cells – first posited by Bretscher and Cohn in 1970 – serves to enforce self-tolerance^1^; B cells are unable to mount productive immune responses if they encounter self-Ag (signal 1) in the absence of T cell co-stimulation (signal 2). Although Ag recognition is not sufficient, it is nevertheless essential for B cells to recruit, engage, and respond to T cell help^2,3^. In addition to driving cell cycle entry and the metabolic remodeling required to sustain an initial round of proliferation, Ag modulates expression of chemokines and chemokine receptors that position Ag-activated B cells in proximity to T cells at the border of their respective zones in secondary lymphoid organs^2,3^. Ag stimulation also upregulates cell surface molecules required for engagement of T cell help, such as ICAM1, CD86 and MHC class II, and primes B cells to respond to CD40L and cytokines (e.g. IL-4) provided by T cells, in part by initiating transcription of *c-Myc*^2,3^. However, if B cells fail to recruit T cell help within a restricted window of time, they either trigger apoptosis, become anergic, or revert to a “naïve-like” state, depending on the strength and duration of Ag stimulation^4,5^. Although the molecular pathways which mediate signal 2 have been extensively explored, the mechanisms which normally restrain Ag-activated B cells and enforce their dependence upon co-stimulation are incompletely understood. Here we identify a role for the NR4A family in this process.

*Nr4a1*-*3* encode a small family of orphan nuclear receptors (NUR77, NURR1, and NOR1 respectively) that are induced by acute and chronic Ag stimulation in lymphocytes, and are thought to function as ligand-independent, constitutively active transcription factors (TFs)^6-10^. In addition, the NR4A family also trigger apoptosis by translocating to the cytosol, binding to BCL2 and inducing a conformational change that exposes its pro-apoptotic BH3-only domain^11,12^. Structural homology and an overlapping expression pattern raise the possibility of functional redundancy among *Nr4a* gene products^6^. Indeed, the *Nr4a* genes are highly upregulated in thymocytes undergoing negative selection, in Tregs, and in anergic or exhausted T cells in mice, where they play a collectively tolerogenic role; the NR4A family are important to mediate thymic negative selection, development and maintenance of Tregs, and the exhaustion and anergy transcriptional programs triggered by chronic Ag stimulation in mature T cells^9,13-20^. Similarly, germline deletion of both *Nr4a1* and *Nr4a3* in mice (but neither one alone) leads to rapid development of a severe myeloproliferative disorder^21^.

We previously characterized a fluorescent reporter of *Nr4a1* expression (NUR77-EGFP BAC Tg) that scales with the extent of Ag stimulation *in vitro* and *in vivo*, and therefore serves to mark naturally occurring autoreactive T and B cells in healthy mice^22-26^. We showed that this reporter correlates with self-reactivity in three different BCR Tg models^25,27-29^. More recently, we have identified a role for NUR77/*Nr4a1* in imposing a novel layer of B cell tolerance *in vivo* by mediating competitive elimination of chronically Ag-stimulated self-reactive B cells in settings with a limiting supply of the B cell survival factor BAFF^28^. We have similarly shown that NUR77 is upregulated by Ag in the highly self-reactive innate-like B1a population where it functions to restrain generation of natural IgM plasma cells^30^. Although expression of *Nr4a* genes is rapidly and robustly upregulated in response to acute BCR stimulation, their function in this context is unknown, and redundancy among the family members has never been explored in B cells.

Here we use germline and conditional mouse models to show that NUR77/*Nr4a1* and NOR1/*Nr4a3* play partially redundant roles to promote apoptosis and to repress proliferation of B cells that receive signal 1 (Ag) in the absence of signal 2 (co-stimulation). By contrast, receipt of co-stimulatory signals bypasses inhibition by the NR4A family. We identify corresponding effects *in vivo* in response to TI-1, TI-2, and TD immunizations. Through unbiased gene expression profiling, we identified a novel set of transcriptional targets of the Nr4a family that are enriched for Ag-induced primary response genes and are cooperatively repressed by NUR77/*Nr4a1* and NOR1/*Nr4a3*, including the transcriptional co-factor BATF which in turn regulates CMYC and B cell proliferation. Unexpectedly, we also identify a role for the NR4A family in restraining B cell access to T cell help by repressing expression of the T cell chemokines CCL3/4, as well as CD86 and ICAM1, and show that this is relevant under conditions of B cell competition for limiting amounts of T cell help. Our work identifies a novel role for the NR4A family as mediators of Ag-dependent negative feedback in B cells. We show that this serves to reinforce tolerance by increasing B cell dependence upon signal 2, and to regulate clonal competition in settings of limiting T cell help.

## MATERIALS AND METHODS

### Mice

NUR77-EGFP and IgHEL (MD4) mice were previously described^25,31^. *Nr4a1*^*fl/fl*^ mice were generously shared by Pierre Chambon and Catherine Hedrick^15^. Mb1-cre, CD21-cre, OTII, *CD40L-/-, Nr4a1-/-*, B1-8i, *Batf-/-*, C57BL/6, and CD45.1+ BoyJ mice were from The Jackson Laboratory ^32-38^. *TCRalpha-/-* were obtained from the Weiss lab^39^. *Nr4a3-/-* mice were generated via electroporation of gRNA and Cas9 mRNA. In brief, Cas9 protein (40μM) and *Nr4a3* gRNAs (80μM) were mixed and electroporated into C57BL/6 zygotes. The sequence of *Nr4a3* exon 3 (containing start ATG) targeted for deletion is shown in Fig 4E. 15 founder lines with the targeted deletion were identified through a screen for PCR amplicon size, and confirmed via sequencing of cloned PCR products. Two founder lines were chosen for further analysis and were backcrossed for at least 4 generations onto the C57BL/6J genetic background. All other strains were fully backcrossed to the C57BL/6J genetic background for at least 6 generations. All mice were housed in a specific pathogen-free facility at UCSF according to University and National Institutes of Health guidelines.

**Figure 1.**
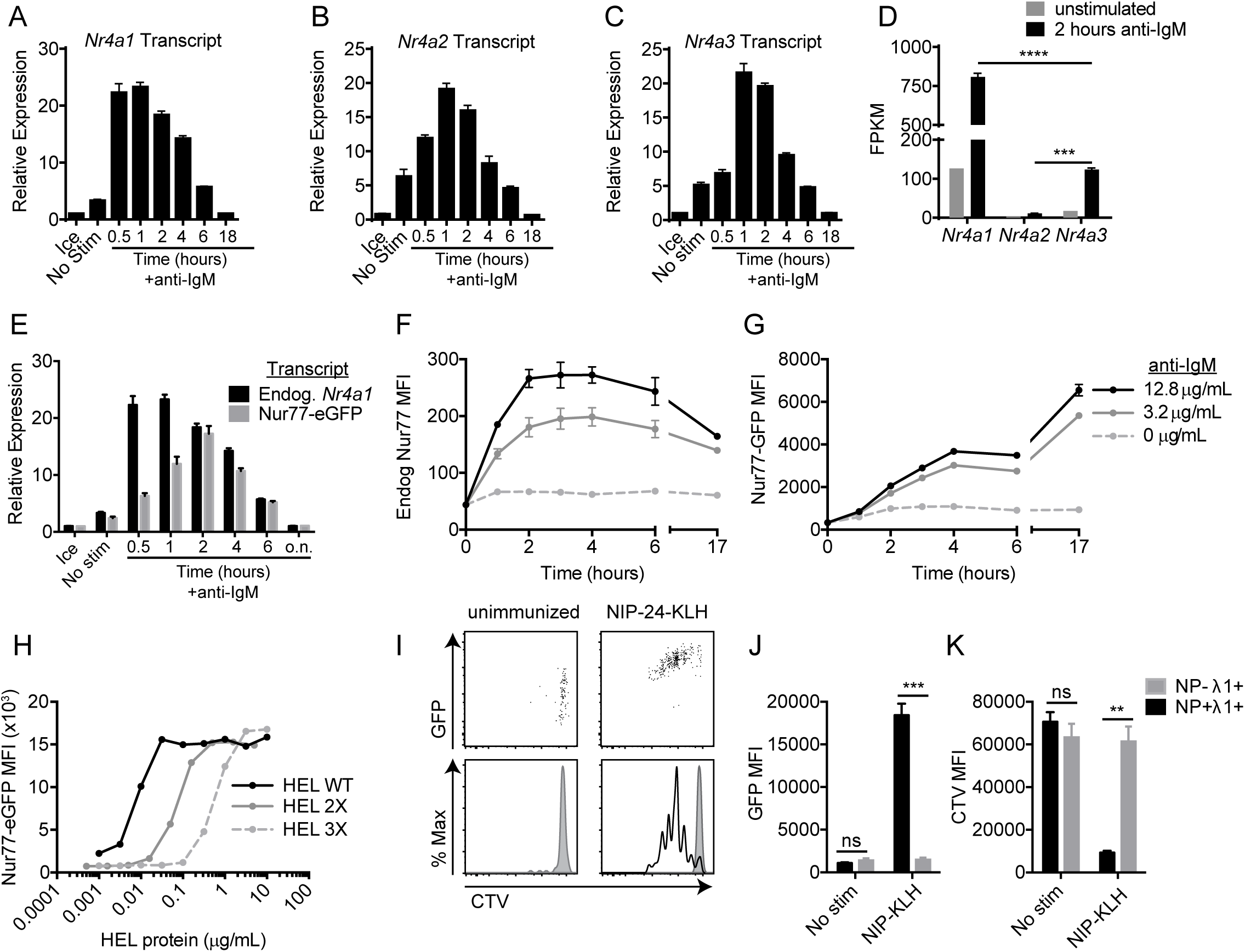
NUR77/Nr4a1 expression scales with Ag stimulation *in vitro* and *in vivo*. **A-C, E**. Lymphocytes from NUR77-EGFP BAC Tg mice were stimulated with 10 μg/mL anti-IgM for the indicated times. qPCR was performed to determine relative expression of *Nr4a1, Nr4a2, Nr4a3*, and EGFP transcript. **D**. B cells from WT mice were purified by bench-top negative selection from pooled splenocytes and LNs, stimulated with 10 μg/mL anti-IgM for 2 h, flash frozen and sent to Q2 Solutions for RNA sequencing. Displayed is the FPKM of *Nr4a1, Nr4a2* and *Nr4a3*. N=1 for the unstimulated conditions, and N=4 for the stimulated conditions. **F-G**. Splenocytes from NUR77-EGFP reporter mice were stimulated with the given doses of anti-IgM for the indicated times. Expression of endogenous NUR77 (F) and EGFP (G) in CD23+ B cells was assessed via flow cytometry. **H**. Lymphocytes from NUR77-EGFP reporter mice were incubated with soluble HEL, HEL 2X, and HEL 3X at serially diluted concentrations for 24 h. Expression of EGFP in B220+ B cells was assessed via flow cytometry. **I-K**. Splenocytes from B1-8i NUR77-EGFP mice were loaded with cell trace violet (CTV). 5×10^6^ cells were adoptively transferred into allotype marked CD45.1+ host mice that were subsequently immunized with 100μg NIP-KLH. Host splenocytes were harvested after 3 days, and CTV, EGFP, NP-binding and λ1 surface expression of donor B cells was assessed via flow cytometry. Data in this figure depict N=3 biological replicates for all panels except (D) as noted above. Mean +/- SEM displayed for all graphs. Statistical significance was assessed with one-way ANOVA with Tukey’s (D); student’s t-test (J, K). ***p<0.001, ****p<0.0001.

**Figure 2.**
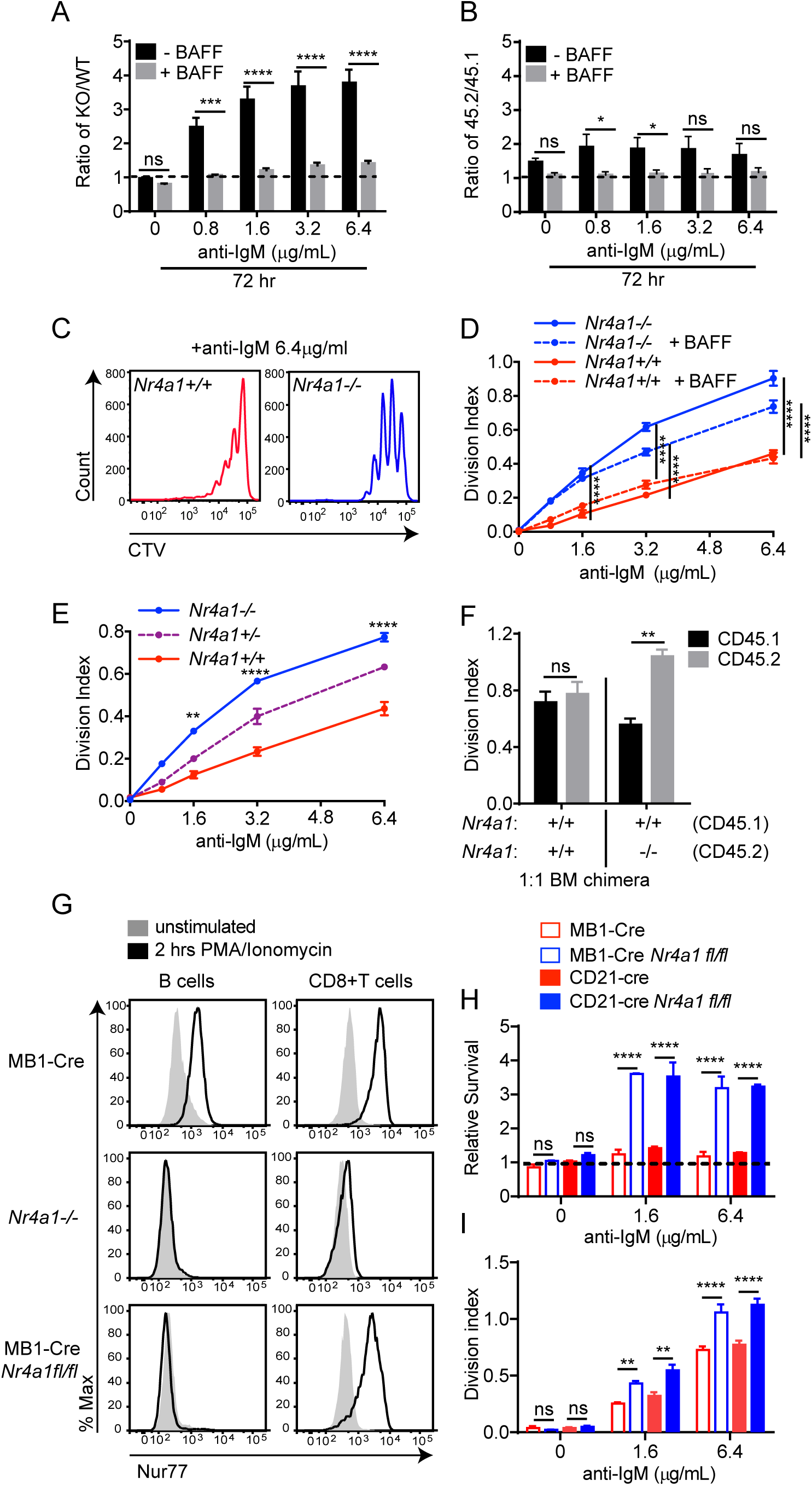
NUR77/*Nr4a1* promotes Ag-induced cell death and restrains Ag-induced proliferation of B cells *in vitro* in a cell-intrinsic manner. **A-D**. Lymphocytes from either *Nr4a1*+/+ or *Nr4a1*-/- CD45.2+ mice were mixed in a 1:1 ratio with CD45.1+ lymphocytes, CTV loaded and co-cultured in the presence of indicated doses of anti-IgM +/- 20 ng/mL BAFF for 72 h. Cells were then stained to detect CD45.1, CD45.2, B220 and CTV via flow cytometry. **A, B**. Displayed is the ratio of CD45.2+ B cells [*Nr4a1-/-* (A) or *Nr4a1*+/+ (B)] relative to co-cultured CD45.1+ WT B cells, normalized to the input ratio. **C**. Representative histograms of CTV dilution in *Nr4a1*+/+ (red) and *Nr4a1*-/- (blue) B cells stimulated with anti-IgM. **D**. Graph depicts division index of co-cultured *Nr4a1*-/- CD45.2+ and *Nr4a1*+/+ CD45.1+ B cells under all conditions assayed. **E**. Lymphocytes from *Nr4a1*+/+, *Nr4a1*+/- and *Nr4a1*-/- mice (littermate progeny of an intercross between *Nr4a1*+/- parents) were CTV loaded and independently cultured in the presence of indicated doses of anti-IgM for 72 h. Cells were then stained to detect B220 and CTV via flow cytometry. Graph depicts division index of each genotype. **F**. Competitive bone marrow chimeras were generated by reconstituting lethally irradiated CD45.1+ WT mice with 1:1 mixtures of either *Nr4a1*+/+ CD45.2+ or *Nr4a1*-/- CD45.2+ bone marrow mixed with WT CD45.1+ bone marrow. 8 weeks after reconstitution, lymphocytes were harvested, loaded with CTV and incubated for 72 h with 5 µg/mL anti-IgM. CTV dilution in B220+ B cells was determined via flow cytometry and graph depicts division index for N=4 biological replicates. **G**. Lymphocytes from mb1-cre, *Nr4a1*-/-, and mb1cre x *Nr4a1* fl/fl mice were stimulated with PMA and ionomycin for 2 hours. Endogenous Nur77 expression was assessed in B220+ B cells and CD8+ T cells by intracellular staining via flow cytometry. Histograms are representative of > 3 biological replicates for each condition. **H-I**. Lymphocytes from mb1-cre, mb1cre x *Nr4a1* fl/fl, CD21-cre, and CD21-cre x *Nr4a1* fl/fl mice were mixed in a 1:1 ratio with WT CD45.1+ lymphocytes, CTV loaded, and co-cultured with the given doses of anti-IgM for 72 h. **H**. Displayed is the ratio of cre+ or cre+ *Nr4a1* fl/fl CD45.2+ B cells relative to co-cultured CD45.1+ WT B cells, normalized to the input ratio. **I**. Graph depicts division index of each genotype. Data in this figure depict N=3 biological replicates for all panels except (F) as noted above. Mean +/- SEM displayed for all graphs. Statistical significance was assessed with student’s t-test with Holm-Sidak (A, B, F, H, I); two-way ANOVA with Tukey’s (D, E). *p<0.05, **p<0.01, ***p<0.001, ****p<0.0001

**Figure 3.**
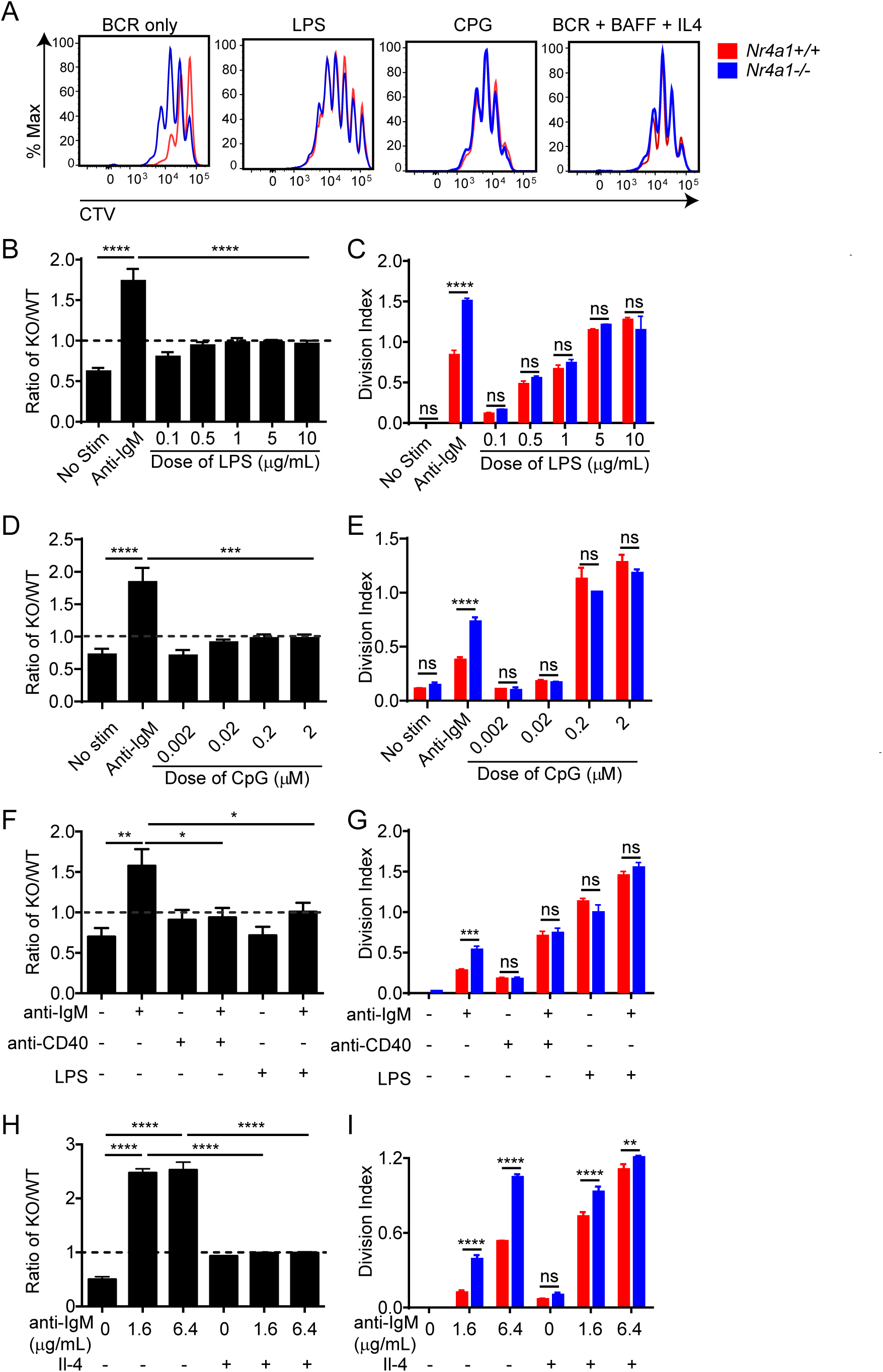
NUR77/*Nr4a1* restrains survival and proliferation of B cells that receive signal one in the absence of signal two. **A-I**. Lymphocytes from CD45.1+ *Nr4a1*+/+ and CD45.2+ *Nr4a1*-/- mice were mixed in a 1:1 ratio and loaded with CTV prior to co-culture for 72 h in the presence of various stimuli as described. Cells were then stained to detect CD45.1, CD45.2, B220 and CTV via flow cytometry. **A**. Histograms depict CTV dilution in B cells stimulated (from left to right) with either 10 μg/mL anti-IgM alone, 5 μg/mL LPS, 0.2 μM CpG, or anti-IgM + 20 ng/mL BAFF + 10 ng/mL IL-4, and are representative of > 3 biological replicates. **B-I**. Doses of stimuli as depicted, except 10 μg/mL anti-IgM (B-G), 1 µg/mL LPS and anti-CD40 (F, G), and 10ng/ml IL-4 (H, I). **B, D, F, H**. Displayed is the ratio of CD45.2+ *Nr4a1*-/- B cells relative to CD45.1+ *Nr4a1*+/+ B cells, normalized to the input ratio. **C, E, G, I**. Graphs depict division index of CD45.1+ *Nr4a1*+/+ (red) and CD45.2+ *Nr4a1*-/- (blue) B cells. Data in this figure depict N=3 biological replicates for all panels. Mean +/- SEM displayed for all graphs. Statistical significance was assessed with one-way ANOVA with Dunnett’s (B, D, F) or Sidak’s (H); student’s t-test with Holm-Sidak (C, E, G, I). Error bars in (B, D) represent comparison of anti-IgM treated condition compared to every other condition. *p<0.05, **p<0.01, ***p<0.001, ****p<0.0001

**Figure 4.**
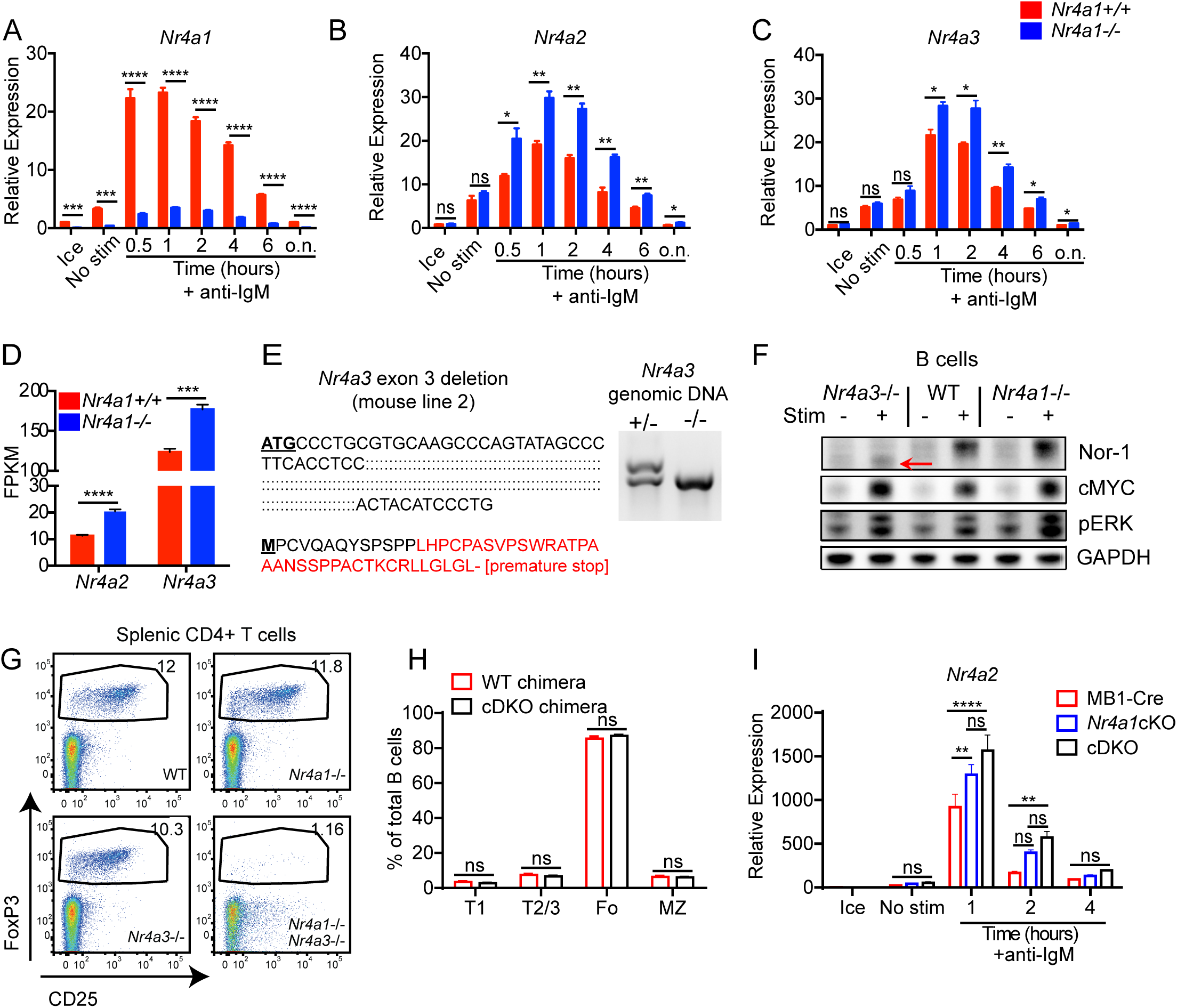
Conditional elimination of NR4A function in B cells. **A-C**. Lymphocytes from *Nr4a1*+/+ and *Nr4a1*-/- mice harboring NUR77-EGFP BAC Tg were stimulated with 10 μg/mL anti-IgM for the indicated times. qPCR was performed to determine relative expression of *Nr4a1, Nr4a2* and *Nr4a3* transcripts. Samples were also subjected to surface staining in parallel to detect % B cells via flow cytometry. Mean % B cells for *Nr4a1+/+* samples was 37.6 ± 0.15 (SEM), and for *Nr4a1*-/- samples was 33.5 ± 1.02 (SEM). N=3 biological replicates for all conditions. *Nr4a1*+/+ samples correspond to data in Figs 1A-C. **D**. Purified B cells from *Nr4a1*+/+ and *Nr4a1*-/- mice were stimulated for 2 h with 10 μg/mL anti-IgM, and analyzed by RNAseq as described in Fig 1D and methods. Displayed is the FPKM of *Nr4a2* and *Nr4a3*. N=1 for the unstimulated condition, and N=4 for the stimulated condition. **E**. Left: Schematic showing the extent of nucleotide deletion in exon 3 of the *Nr4a3* gene harboring ATG translation initiation site, resulting in the introduction of a premature stop codon. Right: Representative PCR showing absence of the WT *Nr4a3* gene and the presence of a truncated product in *Nr4a3*-/- mice. **F**. Purified B cells from WT, *Nr4a1*-/- and *Nr4a3*-/- mice were incubated +/- PMA and ionomycin for 2 h. Subsequently whole cell lysates were blotted with Ab to detect NOR1, c-MYC, pERK, and GAPDH. Red arrow indicates presence of low abundance truncated NOR1 protein. **G**. Representative flow plots showing the gated splenic Treg cell population in WT, *Nr4a1-/-, Nr4a3-/-* and *Nr4a1-/-* x *Nr4a3-/-* mice, as determined by CD25 expression and intra-cellular staining for Foxp3 in splenic CD4+ T cells. Plots are representative of at least 3 WT, *Nr4a1-/-* and *Nr4a3-/-* mice, and 2 DKO mice (*Nr4a1-/- Nr4a3-/-*). **H**. Chimeras were generated by reconstituting lethally irradiated CD45.1+ WT mice with bone marrow from either CD45.2+ mb1-cre mice (cre chimera), or CD45.2+ mb1-cre x *Nr4a1* fl/fl x *Nr4a3-/-* mice (cDKO chimera). 8-10 weeks after reconstitution, mice were sacrificed for analysis by flow cytometry. Shown are the relative sizes of developmental B cell subsets in the spleen as determined via expression of CD21, CD23 and CD93 on gated B cells. N= 5 chimeras / genotype. **I**. Different genotypes are defined as the following: mb1-cre: mb1-cre mice; *Nr4a1*cKO: mb1-cre x *Nr4a1* fl/fl; cDKO: mb1-cre x *Nr4a1* fl/fl x *Nr4a3-/-*. Purified B cells were isolated from whole lymph node via MACs purification, and stimulated for the indicated times with 10 μg/mL anti-IgM. Relative *Nr4a2* transcript was determined via qPCR. Data in this figure depict N=3 biological replicates for all panels except (D) and (H) as noted above. Mean +/- SEM displayed for all graphs. Statistical significance was assessed with student’s t-test with Holm-Sidak (A, B, C); student’s t-test without correction (D, H); two-way ANOVA with Tukey’s (I). *p<0.05, **p<0.01, ***p<0.001, ****p<0.0001

### Antibodies and Reagents

#### Abs for surface markers

Streptavidin (SA) and Abs to B220, CD3, CD19, CD21, CD23, CD86, CD62L, CD69, CD93 (AA4.1), CD44, CD45.1, CD45.2, FAS, GL7, IgM, IgD, Igλ1 conjugated to biotin or fluorophores (Biolegend, eBiosciences, BD, or Tonbo); NP(23)-PE (LGC Biosearch Technologies).

#### Intra-cellular FOXP3 staining

FOXP3/Transcription factor staining buffer set (ebioscience), FOXP3 Ab conjugated to APC (clone FJK-16s) manufactured by Invitrogen.

#### Abs for intra-cellular staining

BATF Ab (clone C7D5), pERK Ab (clone 194g2) and pS6 Ab (clone 2F9), were from Cell Signaling Technologies; NUR77 Ab conjugated to PE (clone 12.14) and EGR2 Ab conjugated to PE (clone erongr2) manufactured by Invitrogen, purchased from eBioscience; c-MYC Ab (clone D84C12, Cell Signaling); activated caspase-3 Ab conjugated to APC (clone C92-605) purchased from BD Pharmingen; IRF4 conjugated to PE (clone 3E4) was from eBioscience; Donkey anti-rabbit secondary Ab conjugated to APC was from Jackson Immunoresearch.

#### Abs for immunoblots

c-MYC Ab (clone D84C12), pERK Ab (Thr202/Tyr204), (clone 197G2), and GAPDH Ab (14C10) were from Cell Signaling; NOR1 Ab (cat# ab94507) was from Abcam; NUR77 Ab (cat# 554088, clone 12.14) was from BD Pharmingen. Anti-mouse- and anti-rabbit-HRP secondary Ab were from Southern Biotech.

#### Stimulatory Abs

Goat anti-mouse IgM F(ab’)2 was from Jackson Immunoresearch; Stimulatory anti-IgD was from MD Biosciences; Murine IL-4 (Peprotech), anti-CD40 (hm40-3 clone; BD Pharmingen), recombinant murine BAFF (R&D), LPS (O26:B6; Sigma), CpG (ODN 1826; InvivoGen), anti-CD40L blocking Ab (clone MR-1, BioXcell).

#### ELISA reagents

Costar Assay Plate, 96 Well Clear, Flat bottom half area high binding, polystyrene (Corning); SA-HRP, anti-mouse IgM, anti-mouse IgG1 and anti-mouse IgG3 conjugated to HRP (Southern Biotech); TMB (Sigma); NP(1)-RSA and NP(29)-BSA (Biosearch); CCL3 and CCL4 ELISAs were performing using the Duoset kit from R&D systems: ancillary reagent kit 2 (cat #:DY008), Mouse CCL3 MIP-1 alpha DuoSet ELISA (R&D #DY450-05), Mouse CCL4 MIP-1 beta DuoSet ELISA (R&D #DY451).

#### Immunogens

NP(53)-Ficoll, NP(17)-OVA, NP(0.3)-LPS and NP(29)-KLH were from Biosearch; Alhydrogel 1% adjuvant (Accurate Chemical and Scientific Corp).

#### Other

Recombinant WT HEL, HEL-2X (R73E, D101R), HEL-3X (R73E, D101R, R21Q) proteins were a gift from Dr. Wei Cheng, University of Michigan^40^. Complete culture media was prepared with RPMI-1640 + L-glutamine (Corning-Gibco), Penicillin Streptomycin L-glutamine (Life Technologies), HEPES buffer [10mM] (Life Technologies), B-Mercaptoethanol [55mM] (Gibco), Sodium Pyruvate [1mM] (Life Technologies), Non-essential Amino acids (Life Technologies), 10% heat inactivated FBS (Omega Scientific).

### Flow Cytometry and data analysis

After staining, cells were analyzed on a Fortessa (Becton Dickinson). Data analysis was performed using FlowJo (v9.9.6 and v10) software (Treestar Inc.). Proliferative indices ‘division index’ and ‘% divided’ were calculated using FlowJo. Statistical analysis and graphs were generated using Prism v6 (GraphPad Software, Inc). Statistical tests used throughout are listed at the end of each figure legend. Student’s unpaired T test was used to calculate p values for all comparisons of two groups, and correction for multiple comparisons across time points or doses was then performed using Holm-Sidak method. One-way or two-way ANOVA with follow-up Tukey’s, Dunnett’s, or Sidak tests were performed when more than two groups were compared to one another. Mean ± SEM is displayed in all graphs. Statistical analysis of RNAseq data is described separately below. Throughout figures: *p<0.05, **p<0.01, ***p<0.001, ****p<0.0001.

### Intracellular staining to detect pERK, pS6, NUR77, cMYC, BATF, EGR2, IRF4, and activated Caspase 3

Either immediately *ex vivo* or following *in vitro* stimulation, cells were fixed in a final concentration of 2% paraformaldehyde for 10 minutes, permeabilized on ice with 100% methanol for 30 minutes (or -20C overnight), and following washes and rehydration were then stained with primary Ab followed by lineage markers and secondary antibodies if needed as previously described ^26^.

### FOXP3 staining

FOXP3 staining was performed utilizing a FOXP3/transcription factor buffer set (eBioscience) in conjunction with an APC anti-FOXP3 Ab (eBioscience), as per manufacturer’s instructions.

### Intracellular Calcium Flux

Cells were loaded with 5 μg/mL Indo-1 AM (Life Technologies) and stained with lineage markers for 15 minutes. Cells were rested at 37°C for 2 minutes, and Indo-1 fluorescence was measured by FACS immediately prior to and after stimulation to determine intracellular calcium.

### Live/dead staining

LIVE/DEAD Fixable Near-IR Dead Cell Stain kit (Invitrogen). Reagent was reconstituted as per manufacturer’s instructions, diluted 1:1000 in PBS, and cells were stained at a concentration of 2×10^6^ cells /100μL on ice for 10 minutes.

### Vital dye loading

Cells were loaded with CellTrace Violet (CTV; Invitrogen) per the manufacturer’s instructions except at 5 ×10^6^ cells/ml rather than 1 × 10^6^ cells/ml.

### In Vitro B Cell Culture and Stimulation

Splenocytes or lymphocytes were harvested into single cell suspension, subjected to red cell lysis using ACK (ammonium chloride potassium) buffer in the case of splenocytes, +/- CTV loading as described above, and plated at a concentration of 2-5 × 10^5^ cells/200 μL in round bottom 96 well plates in complete RPMI media with stimuli for 1-3 days. *In vitro* cultured cells were stained to exclude dead cells as above in addition to surface or intra-cellular markers for analysis by flow cytometry depending on assay.

### Immunoblot analysis

Thymocytes and purified splenic/LN B cells (protocol described below) were harvested from mice, and stimulated in complete media +/- PMA/ionomycin for 2 hours at 37°C. Following stimulation, cells were lysed with 1% NP40, and centrifuged for 15 minutes at 20,000g to remove cellular debris. The supernatants were denatured at 95°C for 5 minutes in SDS sample buffer with 2.5% BME. Lysates were run on Tris-Bis thick gradient (4%-12%) gels (Invitrogen), and transferred to PVDF membranes with a Mini-Protean Tetra cell (Bio-rad). Membranes were blocked for 1 hour with 3% skim milk in TBST, and then probed with primary antibodies listed above, overnight at 4°C. The next day, membranes were incubated with HRP-conjugated secondary antibodies. Blots were developed utilizing a chemiluminescent substrate (Western Lightning Plus ECL, Perkin Elmer), and visualized with a Chemi-doc Touch imager (Bio-rad).

### Immunizations

Mice were immunized via I.P injection of 200 μL of immunogen diluted in PBS. For T-dependent immunizations, NIP(24)-KLH, NP(17)-OVA, or NP(29)-KLH (LGC Biosearch Technologies) either 10 μg or 100 μg was diluted in PBS, and emulsified in Alhydrogel 1% adjuvant (Accurate Chemical and Scientific Corp.). For NP-Ficoll immunization, 100 μg NP(53)-Ficoll (Biosearch) was diluted in PBS. For NP-LPS immunization, 100 μg NP(0.6)-LPS (Biosearch) was diluted in PBS. For GC analysis, mice were sacrificed 7 days after immunization, and spleens were harvested for analysis via flow cytometry. For serum antibody titers, mice were bled prior to immunization, and then serially every seven days for either 21- or 28-days total, and titers were determined via ELISA as described below. Adoptive transfer assays are described further below.

### B cell purification

B cell purification was performed utilizing MACS separation per manufacturer’s instructions. In short, pooled spleen and/or lymph nodes were prepared utilizing the B cell isolation kit (Miltenyi), and purified by negative selection through an LS column (Miltenyi). Processed samples were then subjected to either RNA preparation, immunoblot, *in vitro* culture, or adoptive transfer.

### ELISA for CCL3/CCL4 detection

Supernatants from *in vitro* cultured purified splenic/LN B cells plated at 5×10^5^ cells/200μL were harvested after 48 hours and CCL3/4 concentration was measured in supernatants using commercial ELISA kit per manufacturer’s instructions (R&D biosystems).

### ELISA for serum Ab titer

Serum antibody titers for anti-NP IgM, IgG3 and IgG1 were measured by ELISA. For anti-NP IgM, IgG3, and low affinity anti-NP IgG1 96-well plates (Costar) were coated with 1ug/mL NP(29)-BSA (Biosearch). For high affinity anti-NP IgG1, 96-well plates (Costar) were coated with 1μg/mL NP(1)-BSA (Biosearch). Sera were diluted serially, and total Ag specific titer was detected with a corresponding anti-Ig-HRP (Southern Biotech). All ELISA plates were developed with TMB (Sigma) and stopped with 1N sulfuric acid. Absorbance was measured at 450 nm using spectrophotometer (Spectramax M5, Molecular Devices).

### Bone marrow chimeras

Host mice were irradiated with 530 rads x 2, 4 hours apart, and injected IV with 10^6^ donor A mixed with 10^6^ donor B BM cells to generate competitive chimeras or with 2×10^6^ donor BM cells independently to generate unmixed chimeras. Chimeras were sacrificed 10-12 weeks after irradiation, and cells were assayed ex vivo by flow staining to assess lymphocyte development, or after *in vitro* stimulation as described above to assay protein expression and proliferation as described above.

### Adoptive transfer experiments

In order to study T-dependent responses, splenocytes and lymphocytes from CD45.2 *Nr4a1-/-* B1-8i Tg and CD45.1/2 *Nr4a1+/+* B1-8i Tg mice were harvested into single cell suspensions, subjected to red blood cell lysis with ACK buffer and mixed in a 1:1 ratio. B cells were then purified by negative selection and loaded with CTV vital dye as described above. 2-3×10^6^ cells in 200 μL total volume were injected into each host via the tail vein (+/- varying numbers of OTII splenocytes as noted in figures). One day post-transfer, hosts were either immunized IP with 100μg NP(17)-ova / alum (1:1) or left unimmunized. After 4 days, hosts splenocytes were harvested and analyzed by flow cytometry to detect donor B cells, NP-binding, and CTV dilution. Alternatively, to study T-independent responses (as in Fig 5, S5), B1-8i splenocytes were loaded directly with vital dye and 4-5×10^6^ splenocytes were adoptively transferred into T-sufficient or T-deficient hosts (+/- anti-CD40L blocking Ab). Mice were then immunized IP within 1 hour with either 100ug NIP(24)-KLH (as in Fig 1), 10μg NP(29)-KLH (Fig S5), or 100μg NP(17)-OVA (Fig 5) each mixed 1:1 with alum, and host spleen was harvested for FACS analysis after 3 days.

**Figure 5.**
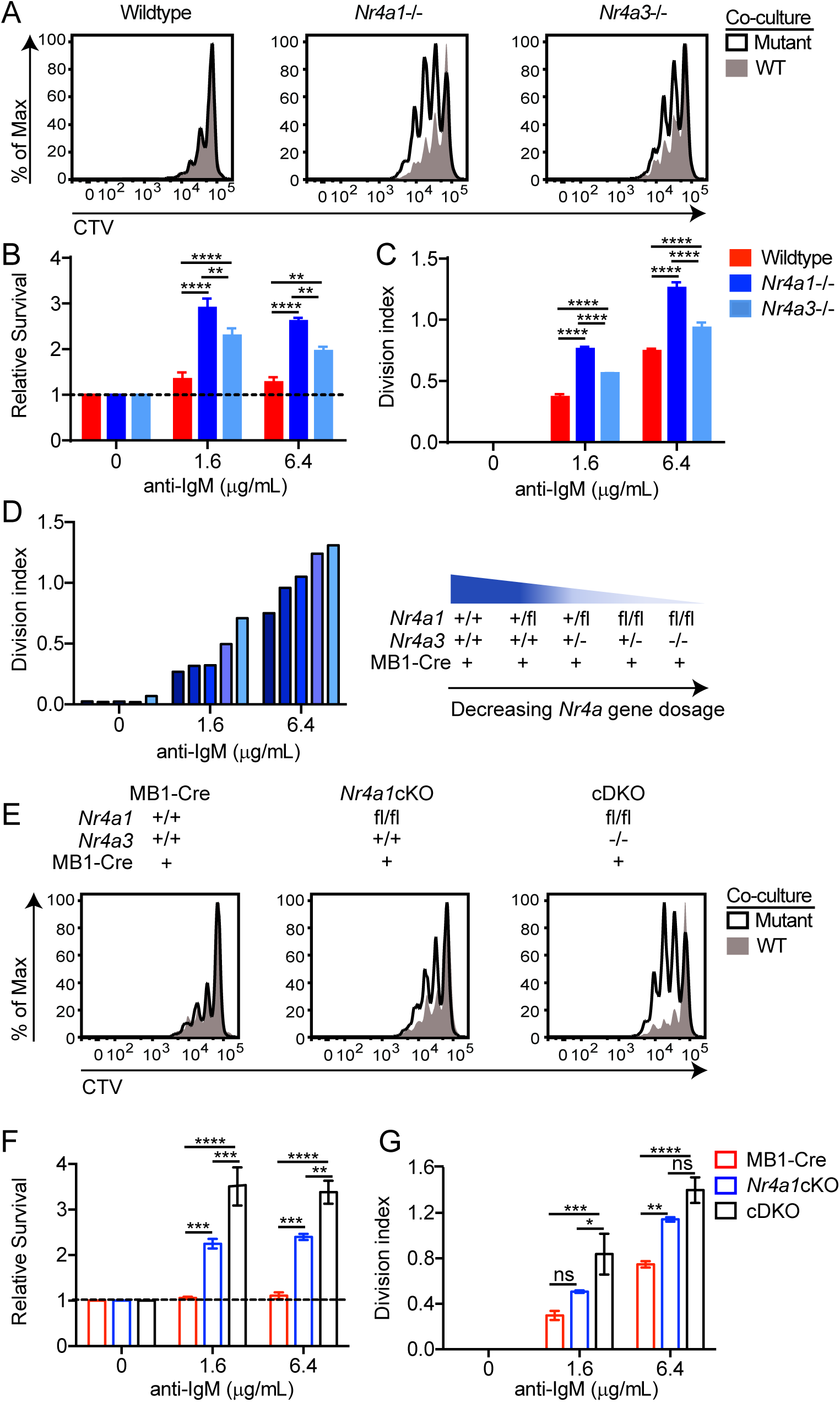
NUR77/*Nr4a1* and NOR1/*Nr4a3* redundantly restrain Ag-induced B cell expansion. **A-C**. Lymphocytes from CD45.2+ WT, *Nr4a1-/-*, and *Nr4a3*-/- mice were each mixed in a 1:1 ratio with CD45.1+ lymphocytes, CTV loaded and co-cultured in the presence of indicated doses of anti-IgM for 72 h. Cells were then stained to detect CD45.1, CD45.2, B220 and CTV via flow cytometry. **A**. Histograms depict CTV dilution in CD45.2+ B cells (black histogram) and co-cultured CD45.1 WT B cells (shaded gray histogram), and are representative of at least 3 mice/genotype. **B**. Shown is the ratio of WT, *Nr4a1-/-* or *Nr4a3-/-* B cells relative to co-cultured CD45.1+ WT B cells, normalized to the unstimulated condition. **C**. Graph depicts division index for each genotype. **D**. An allelic series was generated by crossing *Nr4a3*-/-, mb1-cre and *Nr4a1* fl/fl lines to generate mice with a varying number of functional *NR4A* alleles, as indicated by the legend to the right. Lymphocytes were prepared and co-cultured as described for A-C above. Graph depicts division index plotted for N=1 mouse/genotype and is representative of 3 independent experiments. **E-G**. Different genotypes are defined as the following: mb1-cre: mb1-cre mice; *Nr4a1*cKO: mb1-cre x *Nr4a1* fl/fl; cDKO: mb1-cre x *Nr4a1* fl/fl x *Nr4a3-/-*. Lymphocytes were prepared and co-cultured as described for A-C above. **E**. Histograms depict CTV dilution as described for B above and are representative of at least 3 mice/genotype. **F**. Shown is the ratio of each B cell genotype relative to co-cultured CD45.1+ WT B cells, normalized to the unstimulated condition. **G**. Graph depicts division index for each genotype. Data in this figure depict N=3 biological replicates for all panels except (D) as noted above. Mean +/- SEM displayed for all graphs except (D) as noted above. Statistical significance was assessed with two-way ANOVA with Tukey’s (B, C, F, G). *p<0.05, **p<0.01, ***p<0.001, ****p<0.0001

### qPCR

Either total lymphocytes or MACS-purified B cells were cultured at 37°C with varying stimuli, and harvested into Trizol (Invitrogen), and stored at -80°C. RNA was extracted via phenol phase separation. cDNA was prepared with Superscript III kit (Invitrogen). qPCR reactions were run on a QuantStudio 12K Flex thermal cycler (ABI) using SYBR Green detection (see **Supplementary Table 1** for list of primers).

### RNA sequencing

Single cell suspensions were generated from pooled splenocytes and lymphocytes as described above. B cells were purified via negative selection using MACS kit (Miltenyi Biotech), following the manufacturer’s instructions. B cells were stimulated ± anti-IgM F(ab)’2 (10μg/ml) for two hours, and subsequently pelleted via centrifugation, supernatant was removed, and the pellet was frozen at -80°C. Quadruplicate biological replicate samples of stimulated *Nr4a1+/+* and *Nr4a1-/-* B cells were then sent to Q^2^ lab solutions for RNA preparation, sequencing and analysis. In brief, RNA samples were converted into cDNA libraries with the Illumina TruSeq Stranded mRNA sample preparation kit, and then sequenced on an Illumina sequencing platform. After sequencing, QC analysis and gene and isoform quantification was performed according to the Q2 solutions in-house RNAv9 pipeline. After QC analysis, samples were aligned using STAR software version 2.4 and quantification of FPKM was performed using RSEM version 1.2.14. Genes with FPKM < 10 in WT samples after BCR stimulation were filtered out as were genes with FPKM = 0 in any sample. Differentially expressed genes between WT and KO stimulated biological replicates were then identified on the basis of FPKM. Statistical significance and correction for multiple hypothesis testing was determined with the Holm-Sidak method, with alpha=5.000% using Prism (**Supplemental Data 1**, GEO accession number GSE146747).

## RESULTS

### NUR77/*Nr4a1* expression in B cells scales with Ag stimulation *in vitro* and *in vivo*

*Nr4a1-3* encode a small family of orphan nuclear hormone receptors whose expression, like that of other primary response genes (PRGs), is rapidly induced by a range of mitogenic stimuli^6^. We first sought to characterize their dynamic expression in B cells during response to Ag stimulation. All three transcripts are rapidly but transiently induced by BCR stimulation, as is typical of PRGs (**Figs 1A-C**). However, absolute transcript abundance varies substantially; of the three family members, *Nr4a1* is the most highly expressed in B cells with or without Ag stimulation, *Nr4a3* transcript is approximately 6-fold less abundant than *Nr4a1*, and *Nr4a2* transcript is minimally detectable even after Ag stimulation (FPKM<10) (**Fig 1D**).

We previously characterized a BAC Tg reporter (NUR77-EGFP) in which EGFP is placed under the control of the regulatory region of *Nr4a1*, but does not perturb endogenous *Nr4a1* expression^25^. Both endogenous *Nr4a1* and EGFP transcript are rapidly induced following BCR stimulation. However, EGFP transcript persists longer than endogenous *Nr4a1* transcript, presumably because the reporter lacks a 3’ AU-rich UTR (**Fig 1E**; www.gensat.org). As with *Nr4a1* transcript, endogenous NUR77 protein also exhibits rapid induction and a relatively short half-life, peaking between 2-4 hours after BCR stimulation (**Fig 1F**). Indeed, recent work suggests that NUR77 is ubiquitylated and actively degraded^41^. Although the induction of NUR77-EGFP mirrors that of endogenous NUR77, EGFP protein is not actively degraded, has a relatively long half-life *in vivo* (approximately 20-24 hours), and consequently accumulates over time (**Fig 1G**).

We and others previously showed that NUR77-EGFP expression scales with the intensity of Ag stimulation *in vitro* and *in vivo* in B cells and T cells^19,22-25^. Indeed, both endogenous NUR77 and EGFP expression in reporter B cells scales with the dose of anti-IgM stimulation *in vitro* (**Figs 1F, G**). In order to assess regulation of NUR77 by bona fide Ag, we introduced the IgHEL BCR Tg into the NUR77-EGFP BAC Tg reporter background. We took advantage of this model to titrate both Ag dose and affinity. We stimulated IgHEL Tg B cells with the cognate model Ag HEL (affinity=2×10^10^ M^-1^), as well as HEL derivatives with much lower affinity for the IgHEL BCR (HEL2x: R73E, D101R; affinity = 8×10^7^ M^-1^ and HEL3x: R73E, D101R, R21Q; affinity = 1.5×10^6^ M^-1^)^42^. Induction of NUR77-EGFP scales not only with concentration of Ag, but also with Ag affinity (**Fig 1H)**.

To probe reporter induction *in vivo*, we took advantage of the B1-8i BCR model system in which the VH186.2 heavy chain is knocked into the heavy chain locus, and recognizes the hapten NP when paired with endogenous lambda-1 light chains^43^. In order to track Ag-specific B cells *in vivo*, we generated B1-8i-NUR77-EGFP reporter mice. After adoptive transfer of vital dye-loaded splenocytes from B1-8i reporter mice, congenically marked CD45.1+ hosts were subsequently immunized with NP-conjugated proteins. All detectable NP-binding lambda-1^+^ donor B cells robustly upregulated EGFP and diluted vital dye in response to immunization after 72 hours (**Figs 1I-K**). Conversely, non-NP-binding lambda-1^+^ B cells retained low EGFP expression and did not dilute CTV. These data confirm that EGFP is upregulated in Ag-specific B cells in response to immunization *in vivo*^24^.

### NUR77/*Nr4a1* promotes Ag-induced cell death and restrains Ag-induced proliferation of B cells *in vitro* in a cell-intrinsic manner

We recently identified a role for NUR77/*Nr4a1* in mediating Ag-induced cell death of self-reactive B cells in response to chronic Ag stimulation, but showed that it was dispensable for normal B cell development^28^. Although the NR4A family are also rapidly and transiently upregulated by Ag receptor stimulation in B cells (**Fig 1**), their function in this context nevertheless remains largely unexplored. In order to dissect the role of the NR4A family in acutely Ag-activated B cells, we took advantage of mice with germline deletion of *Nr4a1* since this is the most highly expressed of the three NR4A family members in B cells (**Fig 1D**)^32^. We co-cultured either *Nr4a1-/-* or *Nr4a1+/+* CD45.2+ lymphocytes in a 1:1 ratio with congenically marked CD45.1+ lymphocytes in the presence of increasing doses of anti-IgM. As we previously observed, after 72 hours *Nr4a1*-deficient B cells exhibit a competitive advantage in the presence (but not the absence) of BCR stimulation and this is not attributable to CD45 allotype (**Fig 2A, B**)^28^. Importantly, this is not due to enhanced proximal BCR signal transduction in *Nr4a1-/-* B cells (**Fig S1**). This competitive advantage is suppressed in the presence of the B cell survival factor BAFF, strongly suggesting that it specifically reflects a survival advantage (**Fig 2A**). Of note, BAFF promotes survival of B cells irrespective of Ag stimulation (**Fig S2A**), but does not induce NUR77 reporter expression, and is not independently mitogenic (**Fig S2B**)^44^. We next assessed co-cultured B cells for activated Caspase 3 (a-casp3) at earlier time points to assess initiation of apoptosis. Although a competitive survival advantage is not evident until 48 hours of co-culture, *Nr4a1-/-* B cells exhibit reduced a-casp3 expression relative to WT B cells by 24 hours after BCR stimulation (**Fig S2C, D**). Importantly, both a-casp3 expression and this competitive advantage are suppressed in the presence of BAFF at every time point assayed (**Fig S2E, F**). These data suggest that NUR77 mediates Ag-induced apoptosis of B cells beginning at early time points, and this translates into a competitive survival advantage at later time points.

We observed that NUR77 also restrains proliferation of Ag-stimulated B cells and this phenotype is not rescued by addition of BAFF, suggesting that NUR77 plays distinct and separable roles to promote Ag-induced apoptosis and to restrain proliferation following BCR stimulation (**Fig 2C, D**). This phenotype is *Nr4a1* gene dose-dependent (**Figs S2G, 2E**) and is evident in both mixed and unmixed cultures of either purified B cells or total lymphocytes (**Figs 2D, E, S2H**). Interestingly, as predicted for a negative feedback regulator, loss of NUR77 steepens the dose response curve for Ag-induced B cell expansion (**Fig 2D, E**). IgHEL BCR Tg B cells deficient for *Nr4a1* similarly exhibit both a survival and proliferation advantage in response to BCR ligation, demonstrating that repertoire selection does not account for these phenotypes (**Fig S2I, J**).

To exclude a B cell-extrinsic contribution of NUR77 to these phenotypes, we generated competitive bone marrow chimeras by reconstituting lethally irradiated hosts with a 1:1 mixture of congenically marked CD45.1+ *Nr4a1*^*+/+*^ and CD45.2+ *Nr4a1*^*-/-*^ bone marrow. *Ex vivo* cultures of lymphocytes from these chimeras recapitulate proliferative and survival advantage of *Nr4a1-/-* B cells (**Fig 2F**)^28^. Importantly, control chimeras generated with CD45.1+ and CD45.2+ *Nr4a1+/+* donors exhibited no difference, excluding any contribution of the congenic marker in this assay (**Fig 2F**).

Finally, we took advantage of a conditional allele of *Nr4a1* (distinct from the germline-deficient *Nr4a1* allele used above) to generate mice in which *Nr4a1* was deleted either early or late during B cell development, with mb1-cre or CD21-cre respectively (**Fig 2G**)^15,33,35^. Competitive co-cultures of mb1-cre *Nr4a1*^*fl/fl*^, CD21-cre *Nr4a1*^*fl/fl*^ lymphocytes, or cre controls each mixed with congenically marked CD45.1+ lymphocytes also recapitulate the survival and proliferative advantage of *Nr4a1*-deficient B cells (**Figs 2H, I, S2K, L**). This is comparable to the advantage we observed in B cells from *Nr4a1-/-* mice harboring germline deficiency of NUR77, suggesting that this phenotype is not due to a role for *Nr4a1* either early during B cell development, or in other cell types, nor is it attributable to allele-specific effects (**Fig S2M, N**).

### NUR77/*Nr4a1* selectively restrains *in vitro* survival and proliferation of B cells that receive signal one in the absence of signal two

Activation of B cells by BCR stimulation alone (signal one) typically leads to an abortive round of proliferation followed by apoptosis^5^. In contrast, co-stimulatory signals from T cells or TLR ligands can rescue Ag-stimulated B cells and support humoral immune responses including class switch recombination and plasma cell differentiation^2^. Therefore, we next sought to determine how specific co-stimulatory signals regulate NUR77 expression in B cells. We previously reported that neither BAFF nor IL-4 could induce NUR77-EGFP expression, while both CD40 and the TLR4 ligand LPS could do so – albeit less efficiently than BCR stimulation, suggesting the reporter is sensitive to canonical NF-κB signaling (**Fig S3A-C**)^25,26^. We next sought to define the kinetics of endogenous NUR77 expression in response to these stimuli both in isolation and when combined with BCR ligation (as might be the case *in vivo*). As with Ag receptor stimulation alone, we found that NUR77 expression peaked at 2-4 hours irrespective of co-stimulatory input, implying that the duration of NUR77 expression was not altered by such inputs (**Fig 1F, S3A-C**).

TLR ligands can serve as mitogenic stimuli for B cells either in isolation or together with Ag stimulation^2^. We first titrated doses of either LPS or CpG *in vitro* and found no survival or proliferative advantage for *Nr4a1-/-* B cells at any dose across a very broad titration (**Figs 3A-E, S3D**). The TLR9 ligand CpG mimics viral nucleic acids that are encountered by B cells typically in the context of an intact virion and synergistically drives B cell activation in conjunction with Ag stimulation^45^. Therefore, we titrated CpG dose in the context of BCR stimulation and found that *Nr4a1-/-* B cells lose their advantage with high doses of CpG superimposed on anti-IgM (**Fig S3E, F**). Similarly, co-stimulation of B cells with LPS and anti-IgM eliminated the advantage enjoyed by *Nr4a1-/-* B cells stimulated through BCR alone (**Fig 3F, G**).

T cells deliver essential co-stimulatory signals to B cells, exemplified by CD40L and IL-4, that synergize with Ag to promote B cell survival and proliferation. We found that both anti-CD40 and IL-4, when added to anti-IgM, similarly eliminate the competitive advantage of BCR-activated *Nr4a1-/-* B cells (**Fig 3F-I**). While TLR co-stimulation is usually delivered in conjunction with Ag stimulation (e.g. bacteria, virus), B cells typically experience a physiologic time delay *in vivo* between Ag encounter and recruitment of T cell help. In order to mimic this delay, we systematically varied the time between initial BCR ligation and addition of anti-CD40. We found that addition of signal 2 at early time points eliminated the advantage of *Nr4a1-/-* B cells, but this advantage was again evident if provision of signal 2 was substantially delayed (**Fig S3G**). These data collectively suggest that NUR77 selectively restrains survival and expansion of B cells that receive signal 1 (Ag) alone, and may help to render B cells dependent upon rapid receipt of signal 2 within a fixed time window following Ag encounter.

### Compensation and redundancy among the NR4A family in B cells

Although our studies identify non-redundant functions of NUR77 in B cells (**Figs 2, 3**), the literature provides compelling evidence for profound functional redundancy among the NR4A family in other immune cells, most strikingly in Tregs^13-16,21^. Therefore, we next sought to assess compensatory expression and redundant functions among the NR4A family members in B cells. *Nr4a1-3* are all induced by BCR stimulation, and show similar kinetics of induction and decay (**Fig 1A-C**). Importantly, in the absence of NUR77 expression, *Nr4a2* and *Nr4a*3 exhibit modestly enhanced induction in response to BCR stimulation, as detected both by qPCR and RNAseq, suggesting that they may be negatively regulated by NUR77 (**Fig 4A-D**). NUR77 may also repress its own expression; we generated *Nr4a1-/-* NUR77-EGFP BAC Tg mice and observed enhanced EGFP transcript and protein induction in response to BCR stimulation in the absence of endogenous NUR77 expression (**Fig S4A, B**).

We hypothesized that loss of *Nr4a1* and *Nr4a3* gene products (NUR77 and NOR1 respectively) should eliminate virtually all NR4A function in B cells since *Nr4a2* transcript is minimally expressed in B cells (**Fig 1D**). Importantly, although *Nr4a2* is modestly upregulated in *Nr4a1-/-* B cells, absolute transcript abundance remains quite low relative to *Nr4a3*, even after BCR stimulation (**Fig 4D**). To begin to test this hypothesis, we first generated a novel germline deletion of *Nr4a3* using CRISPR-mediated NHEJ on the *Nr4a1*^*fl/fl*^ genetic background. We electroporated fertilized zygotes with recombinant Cas9 and sgRNAs targeting exon 3 (containing translation start codon ATG) of *Nr4a3*. 15 independent founders were generated, 2 of which were selected for further breeding and study. These lines exhibited a 115 bp and 112bp deletion respectively that resulted in an identical frameshift and premature stop codon (**Fig 4E**).

CRISPR-mediated knockout mice have recently been reported to generate truncated protein products lacking n-terminus, due in part to translation initiation from downstream internal start sites^46^. We therefore sought to confirm that our novel *Nr4a3* mutant represented a loss-of-function allele. Although *Nr4a3* transcript abundance is unaltered and evades nonsense-mediated decay in *Nr4a3-/-* B cells (**Fig S4C, D**), little protein is produced; residual truncated protein expression - detected using a polyclonal Ab raised against aa200-300 of NOR1 (ab94507) - was minimally inducible and of extremely low abundance (**Fig 4F, S4E**). Together this is consistent with highly inefficient initiation of translation from an internal ATG that is downstream of the premature stop codon generated by CRISPR-mediated deletion.

To further confirm that this is indeed a complete loss-of-function allele, we generated germline DKO mice lacking both *Nr4a1* and *Nr4a3*, and observed early development of a Scurfy-like disease characterized by growth arrest, skin lesions, enlarged lymph nodes, and a near-complete lack of peripheral Tregs, as previously described for CD4-cre *Nr4a1*^*fl/fl*^ combined with an independently generated *Nr4a3-/-* line (**Fig 4G**, DNS)^15^. These data collectively demonstrate that our newly generated *Nr4a3-/-* mice harbor a bona fide germline loss of function allele.

We postulated that by combining the mb1-cre *Nr4a1*^*fl/fl*^ model with our new germline *Nr4a3-/-* mice (herein referred to as cDKO), we could eliminate NR4A function in B cells without provoking systemic inflammatory disease associated with deletion of multiple NR4A family members in the germline or in the T cell compartment (**Fig 4G**)^15,21^. We first established that T cell subsets and homeostasis as well as B cell development were unperturbed in *Nr4a3-/-* animals (**Figs S4F-K**). Similarly, T cell homeostasis was unaffected in cDKO mice (**Fig S4L, M**). Further, B cell subsets were not grossly altered in cDKO mice, either in 100% chimeras (**Fig 4H**) or in competitive chimeras (**Fig 4SN, O**), suggesting that the NR4A family is dispensable for development and maintenance of B cells under steady state conditions. Importantly, we identify only modest over-induction of *Nr4a2* in cDKO B cells, suggesting that *Nr4a2* expression is unlikely to compensate for loss of *Nr4a1* and *Nr4a3* in the B cell compartment (**Fig 4I**).

### NUR77/*Nr4a1* and NOR1/*Nr4a3* collectively restrain Ag-induced B cell expansion *in vitro*

We next sought to define whether B cells from *Nr4a3-/-* mice partially or completely phenocopy *Nr4a1-/-* B cell responses to Ag *in vitro*. We found that *Nr4a3-/-* B cells also displayed a competitive survival and proliferative advantage relative to co-cultured wild type B cells, but it was not as pronounced as the advantage enjoyed by *Nr4a1-/-* B cells in parallel assays (**Figs 5A-C, S5A, B**).

We showed that *Nr4a3* expression is over-induced in *Nr4a1-/- B* cells after Ag encounter (**Fig 4C, D**). More importantly, *Nr4a1* and *Nr4a3* play redundant roles in other immune cell types^13-16,21^. Therefore, we next sought to determine whether this was also the case in Ag-stimulated B cells. To do so, we took advantage of mb1-cre *Nr4a1*^*fl/fl*^ *(Nr4a1* cKO forthwith) and *Nr4a3-/- mice* in order to generate an ‘allelic series’ of mice with varying numbers of functional *NR4A* alleles in the B cell compartment (**Fig 5D-G**). We found that Ag-induced survival and proliferation of B cells was enhanced in proportion to the number of deleted *NR4A* alleles (**Figs 5D-G, S5C-D)**.

Because *Nr4a3* is deleted in the germline in cDKO mice, we wanted to rule out cell-extrinsic contributions to these phenotypes. To do so, we generated competitive chimeras and showed that cDKO B cells indeed exhibit a B cell-intrinsic survival and proliferative advantage relative to control B cells in response to Ag stimulation (**Fig S5E-H**). As we observed for *Nr4a1-/-* B cells (**Fig 3**), this advantage for cDKO B cells is lost with the addition of signal 2 (**Fig S5I, J**). Importantly, control competitive chimeras show no such advantage for cre-only B cells (**Fig S5E-J**). We conclude that *Nr4a1* and *Nr4a3* play partially redundant roles in restraining survival and proliferation of B cells in response to BCR stimulation *in vitro*.

### NUR77/*Nr4a1* restrains humoral responses to Ag in the absence of co-stimulation *in vivo*

Next, to test *in vivo* relevance of these *in vitro* observations, we probed immune response to a T-independent BCR stimulus, NP-Ficoll. We previously reported enhanced IgM responses to NP-Ficoll immunization of *Nr4a1-/-* hosts and here observe profoundly increased IgG3 responses to this immunogen across multiple time points (**Fig 6A**)^30^. NP-Ficoll responses are completely dependent upon intact BCR signal transduction^47^. We next sought to determine whether provision of signal 2 in the form of either TLR ligand or engagement of T cell help could eliminate the advantage of *Nr4a1-/-* mice. Indeed, total NP-specific titers induced by NP-LPS or the T-dependent immunogen NP-KLH are comparable irrespective of NUR77 expression (**Figs 6B-C, S6A**). Affinity maturation in response to NP-KLH is also unaffected (**Fig 6D**).

**Figure 6.**
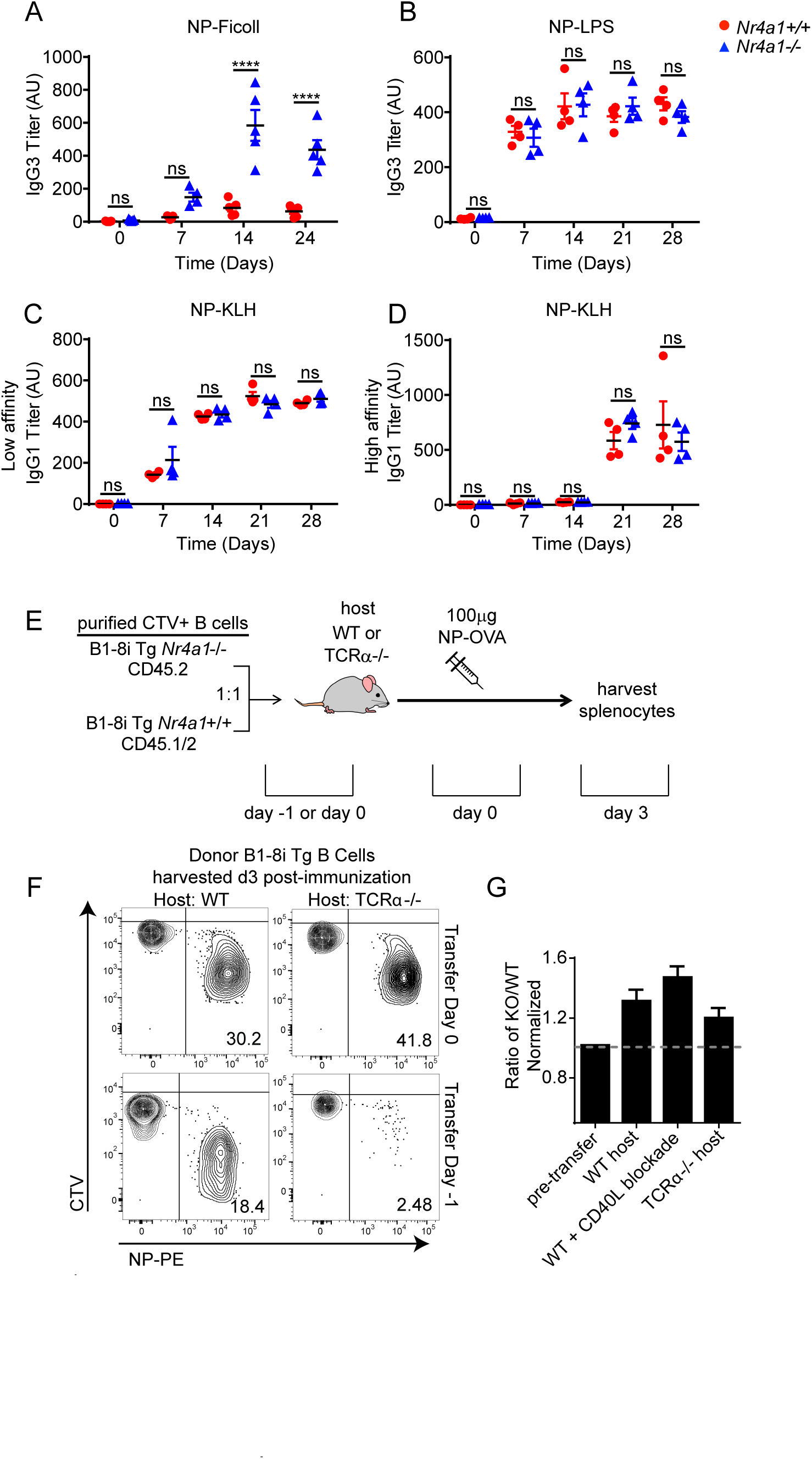
NUR77/*Nr4a1* selectively restrains humoral immune responses to T-independent II immunogens *in vivo*. **A, B**. *Nr4a1*+/+ and *Nr4a1*-/- mice were immunized IP with either 100 μg NP-Ficoll (A) or 100 μg NP-LPS (B) and anti-NP IgG3 titers were determined via ELISA at serial time points. **C, D**. *Nr4a1*+/+ and *Nr4a1*-/- mice were immunized IP with 100 μg NP-KLH admixed 1:1 with alum and anti-NP IgG1 were determined via ELISA at serial time points. ELISA plates were coated with either NP-24-BSA (C), or NP-1-RSA (D) in order to detect total and high affinity NP-specific Abs respectively. **E-G**. Diagram of experimental design (E) for adoptive transfer assays depicted in (F, G). B cells were purified from splenocytes harvested from *Nr4a1*+/+ B1-8 Tg CD45.1/2+ and *Nr4a1*-/- B1-8 Tg CD45.2+ mice via bench-top negative selection, mixed 1:1, and loaded with CTV. 5×10^6^ cells were adoptively transferred into WT or TCRα-/- host mice either 1 day prior to or same day as 100 µg NP-OVA/alum IP immunization, followed by spleen harvest on d3. **F**. Representative flow plots gated on donor B cells depict CTV dilution among NP-binding B cell under varied conditions. **G**. Adoptive transfers performed as depicted in (E) except all hosts were immunized on day of transfer (day 0 condition) and a cohort of WT recipients also received 600 μg of anti-CD40L antibody at the time of adoptive transfer. Shown is the ratio of *Nr4a1*-/- to *Nr4a1+/+* donor B cells among NP+ population normalized to the same ratio in the NP-population. Statistical analyses of these data are depicted in Fig S6B. Data in A-D depict N=4-5 mice /genotype, and data in G depicts N=4 recipient mice per condition. Data in G depict N=4. Data in A, C, D are representative of 2-3 independent experiments. Mean +/- SEM displayed for all graphs. Statistical significance was assessed with student’s t-test with Holm-Sidak (A-D). ****p<0.0001

We next took an adoptive transfer approach in order to isolate the B cell-intrinsic role of NUR77 *in vivo* at early time points after Ag engagement. To do so, we co-transferred congenically marked NP-specific B cells from *Nr4a1*-sufficient or -deficient donors into either T cell-sufficient or -deficient hosts (**Fig 6E**). We sought to develop an *in vivo* assay in which T-independent responses by donor B cells could be tracked. We therefore opted to immunize hosts with a T-dependent immunogen (NP-OVA or NP-KLH) immediately after adoptive transfer, but before B cells could home to the B cell zone in lymphoid follicles, and position themselves for rapid access to cognate T cell help. Indeed, we confirmed, using T-deficient hosts, that such a strategy produced robust T-independent Ag-specific B cell proliferation (**Fig 6F**). By contrast, delay of immunization by 24 hours resulted in a highly T-dependent B cell response (**Fig 6F**). In response to immediate immunization, NP-specific *Nr4a1-/-* B cells exhibit a competitive advantage at early time points (**Figs 6G, S6B, C**). We used two independent strategies (*TCRalpha-/-* hosts and CD40L blockade) to confirm that this advantage was indeed T-independent (**Fig 6G, S6B**). Collectively, these data suggest that, similar to *in vitro* assays (**Figs 2, 3, 5)**, NUR77 selectively restrains B cell responses to Ag in the absence of signal 2 and does so in a B cell-intrinsic manner *in vivo* (see model, **Fig S6D**).

### NUR77/*Nr4a1* and NOR1/*Nr4a3* cooperate to restrain expression of a subset of Ag-induced target genes

Since BCR signal transduction is unaltered in *Nr4a1*^*-/-*^ B cells (**Fig S1**), we next sought to define transcriptional targets of NUR77 that might account for its negative regulatory role downstream of Ag stimulation in B cells. To do so, we assessed global transcript abundance via RNA-Seq in *Nr4a1*^*-/-*^ and *Nr4a1*^*+/+*^ B cells stimulated with anti-IgM for 2 hours (**Fig 7A**). We sought to capture the genes regulated by NUR77 at its peak of expression (2-4 hours) and to enrich for direct targets of NUR77 by selecting such an early time point. A substantial fraction (approx. 14%) of the naïve B cell transcriptome is rapidly upregulated in response to Ag stimulation, as has been previously described (**Fig 7B**)^3,48^. We found a relatively small number of genes that were differentially expressed (DEGs) between *Nr4a1*^*-/-*^ and *Nr4a1*^*+/+*^ B cells after two hours of BCR stimulation (**Fig 7B, Supplementary Data 1**). Genes that were over-induced in *Nr4a1-/-* B cells were highly enriched for Ag-induced PRGs (51%; **Fig 7B**). Reassuringly, this set of genes included *Nr4a2* and *Nr4a3* (**Figs 4B-D**). By contrast, we observed no enrichment for PRGs among genes that were downregulated in *Nr4a1-/-* B cells (**Supplementary Data 1**). This suggests that NUR77 imposes negative feedback regulation downstream of Ag encounter to restrain expression of a subset of PRGs, and we focused our attention on these genes.

**Figure 7.**
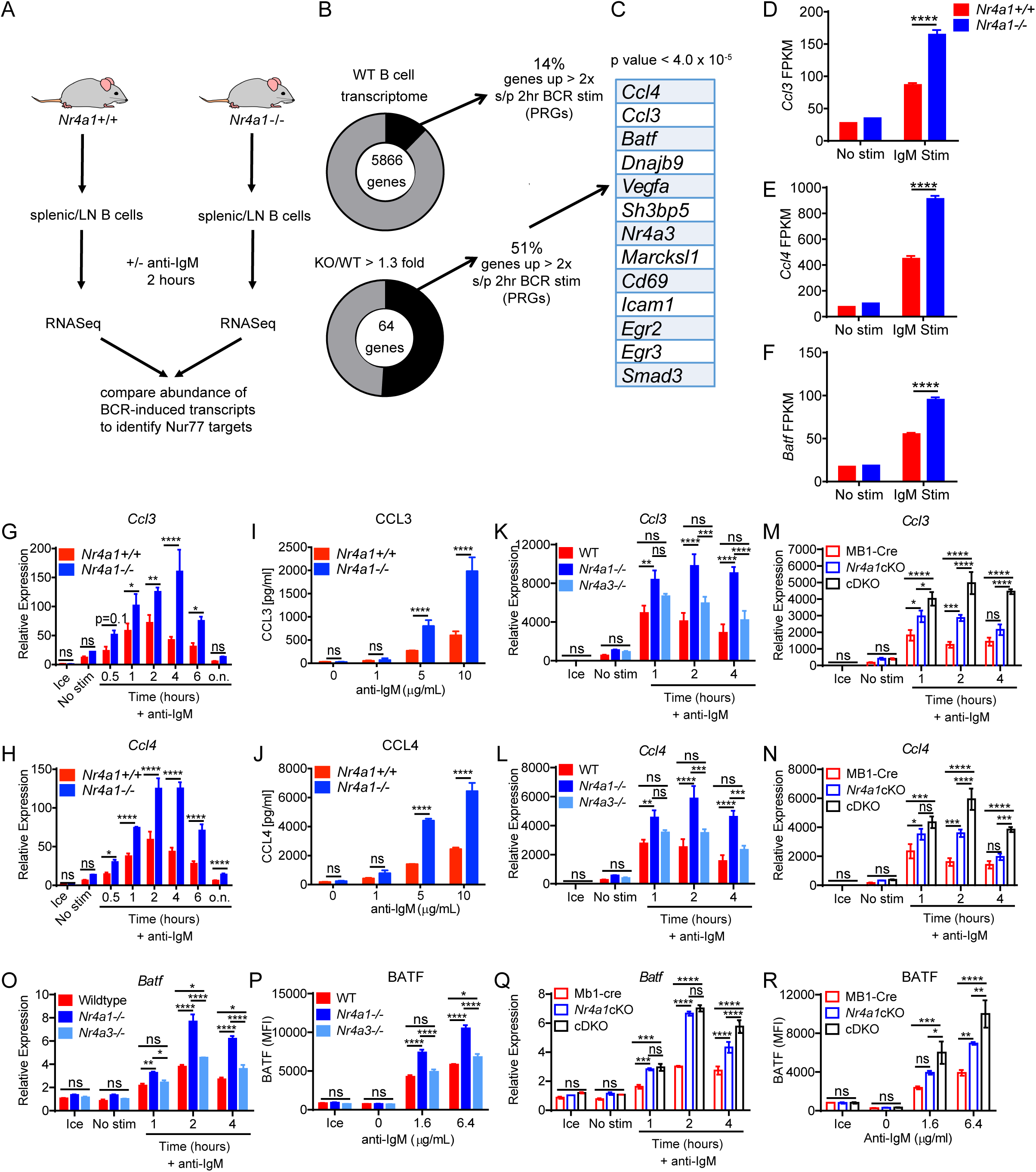
NUR77/*Nr4a1* restrains a subset of Ag-induced primary response genes. **A-F**. B cells from *Nr4a1*+/+ and *Nr4a1-/-* mice were purified by bench-top negative selection from pooled splenocytes and LNs, stimulated with 10 μg/mL anti-IgM for 2 h, flash frozen and sent to Q2 Solutions for RNA sequencing. N=1 for the unstimulated conditions, and N=4 for the stimulated conditions. **A**. Schematic of experimental design. **B**. Graph depicts proportion of genes that are induced > 2x after BCR stimulation in WT B cells (top) and in enrichment in this proportion among all genes with mean FC > 1.3 in KO relative to WT (bottom). **C**. Shown from most to least over-induced are all genes and >2x induced by BCR stimulation with FC > 1.3 in KO/WT (p < 4.0×10^−5^). **D-F**. Displayed is the FPKM of *Ccl3, Ccl4*, and *Batf*. **G-H**. Lymphocytes from *Nr4a1*+/+ and *Nr4a1*-/- mice harboring NUR77-EGFP BAC Tg were stimulated with 10 μg/mL anti-IgM for the indicated times. qPCR was performed to determine relative expression of *Ccl3* (G) and *Ccl4* (H) transcripts. Samples correspond to those described in Fig 4A-C. **I-J**. B cells from *Nr4a1*+/+ and *Nr4a1-/-* mice were purified by bench-top negative selection from pooled splenocytes and LNs, stimulated with given doses of anti-IgM for 48 h, and supernatant was analysed via ELISA to determine concentration of CCL3 (I) and CCL4 (J). Lymphocytes were harvested from *Nr4a1*-/- and *Nr4a1*+/+ mice, and stimulated with the given dose of anti-IgM. The supernatant from the culture was collected at 48 hours, and the concentration of either CCL3 (**I**) or CCL4 (**J**) protein was determined via ELISA. **K, L**. B cells from *Nr4a1*+/+, *Nr4a1-/-*, and *Nr4a3-/-* mice were purified by bench-top negative selection from pooled splenocytes, and were stimulated with 10 μg/mL anti-IgM for the indicated times. qPCR was performed to determine relative expression of *Ccl3* (K) and *Ccl4* (L) transcripts. **M, N**. Experiment performed as in K, L except with purified B cells from genotypes: mb1-cre; mb1-cre x *Nr4a1* fl/fl (“*Nr4a1*cKO”); mb1-cre x *Nr4a1* fl/fl x *Nr4a3-/-* (“cDKO”). qPCR was performed to determine relative expression of *Ccl3* (M) and *Ccl4* (N) transcripts. **O**. Purified, stimulated B cell samples described in K, L subjected to qPCR to determine relative expression of *Batf* transcript. **P**. Lymphocytes were harvested from WT, *Nr4a1-/-*, and *Nr4a3-/-* mice, mixed in a 1:1 ratio with CD45.1 WT lymphocytes, and stimulated with the given doses of anti-IgM for 24 hours. Graph depicts MFI of intracellular BATF protein expression in CD45.2 B cells as determined via flow cytometry. **Q**. Purified, stimulated B cell samples described in M, N subjected to qPCR to determine relative expression of *Batf* transcript. **R**. Experiment performed as in (P) except with genotypes listed. Graph depicts MFI of intracellular BATF protein expression in B cells as determined via flow cytometry. Data in this figure depict N=3 biological replicates for all panels except (A-F) as noted above. Mean +/- SEM displayed for all graphs. Statistical significance was assessed with student’s t-test with Holm-Sidak (D-J); two-way ANOVA with Tukey’s (K-R). *p<0.05, **p<0.01, ***p<0.001, ****p<0.0001

To further prioritize this list of putative NUR77 target genes for follow-up, we filtered DEGs on the basis of statistical significance, which served to identify a small number of genes, many of which are known to play important roles in mediating humoral immune responses (**Fig 7C**)^49-57^. Of these, the most differentially induced genes we identified encoded the T cell chemokines CCL3 and CCL4 (also known as MIP-1 alpha and beta), and the transcription factor BATF (**Figs 7D-F**)^55-57^. *Ccl3* and *Ccl4* were previously shown to be repressed by NR4A over-expression in LPS-stimulated macrophage cell lines^58^. To validate these putative target genes in B cells, we assessed transcript expression by qPCR in BCR-stimulated *Nr4a1*^*-/-*^ and *Nr4a1*^*+/+*^ B cells at serial time points. We found that expression of *Ccl3* and *Ccl4* was rapidly and transiently induced in WT B cells, but both the peak and duration of transcript expression was increased markedly in the absence of NUR77 (**7G, H**). This in turn correlated with increased CCL3 and CCL4 chemokine secretion by *Nr4a1*^*-/-*^ B cells after 48 hours of *in vitro* BCR stimulation (**Fig 7I, J**). By contrast, *Nr4a3-/-* B cells exhibited minimal change in *Ccl3* and *Ccl4* transcript induction (**Fig 7K, L**). Strikingly, however, cDKO B cells that were deficient for both *Nr4a1* and *Nr4a3* expression showed a substantial increase in *Ccl3* and *Ccl4* expression relative to mb1-cre *Nr4a1*^*fl/fl*^ *(Nr4a1* cKO) B cells, suggesting that NUR77/*Nr4a1* and NOR1/*Nr4a3* cooperatively restrain expression of these target genes in B cells (**Fig 7M, N**). Since conditional mutant animals recapitulate *Ccl3/4* over-induction observed in the *Nr4a1-/-* germline-deficient model, we further conclude that these effects are B cell intrinsic.

We took an analogous approach to validate additional putative NR4A target genes; we found that *Batf* transcript and protein were over-induced in *Nr4a1-/-* B cells but not in *Nr4a3-/-* B cells (**Figs 7O, P S7A, B**). Similar to what we found for *Ccl3/4, Batf* transcript was over-induced in cDKO B cells relative to *Nr4a1* cKO B cells at later time points, and this corresponded to over-expression of BATF protein after 24 hours of BCR stimulation (**Fig 7Q, R**). Importantly, we showed that this was reproducible in competitive chimeras as well, confirming B cell intrinsic regulation by the NR4A family (**Fig S7C**). We also went on to validate *Cd69* and *Egr2* as NR4A target genes. Similar to *Batf*, both *Cd69* and *Egr2* transcript and protein products are over-induced in *Nr4a1-/-* B cells but not *Nr4a3-/-* B cells, while cDKO B cells exhibit synergistic and cell-intrinsic over-induction (**Figs S7D-O**). Although it has been suggested that *Irf4* is a target of NUR77 in CD8 T cells, we find only subtle differences in B cells for *Irf4* transcript at early time points and protein at later time points (**Supplementary Data 1, Fig S7P**)^59^. Collectively, these data support a model in which *Nr4a1* and *Nr4a3* are rapidly induced by Ag stimulation and feedback to cooperatively restrain the expression of a small subset of other PRGs that play important roles in orchestrating humoral immune responses (see model, **Fig S7Q**).

### BATF mediates negative regulation of cMYC expression and proliferation by NUR77/*Nr4a1*

We next wanted to understand how NUR77 transcriptional targets could account for enhanced Ag-induced B cell expansion in its absence. Not only do *Nr4a1-/-* B cells proliferate more in response to BCR stimulation (**Fig 2**), but cell size is markedly increased relative to WT after 24 hours of stimulation. (**Fig 8A**). c-MYC protein levels have been shown to correlate with and direct cell growth, metabolic reprogramming, and proliferative potential of naïve and GC B cells in a dose-dependent manner^60-62^. *c-Myc* has also been proposed as a putative NR4A target gene in myeloid cells^63^. Indeed, we find that c-MYC protein expression is markedly over-induced in BCR-stimulated *Nr4a1-/-* and cDKO (but not *Nr4a3-/-*) B cells in a cell-intrinsic manner (**Fig 8B, S8A-C**). Moreover, c-MYC protein is robustly upregulated in both *Nr4a1-/-* and *Nr4a1+/+* B cells with addition of IL-4, and this could account (in whole or in part) for how provision of signal 2 might bypass negative regulation by NUR77 (**Fig 8B**). Therefore, we hypothesized that negative regulation of c-MYC contributes to NR4A-dependent repression of B cell proliferation in response to signal 1 received in the absence of signal 2. However, we only observed a subtle over-induction of *c-Myc* transcript in *Nr4a1-/-* B cells after BCR stimulation (**Fig 8C**), and c-MYC protein over-induction is only detectable at relatively late time points (24 hrs) after Ag stimulation of *Nr4a1-/-* B cells (**Figs 8B, D, E**), suggesting that it may be an indirect transcriptional target of the NR4A family.

**Figure 8.**
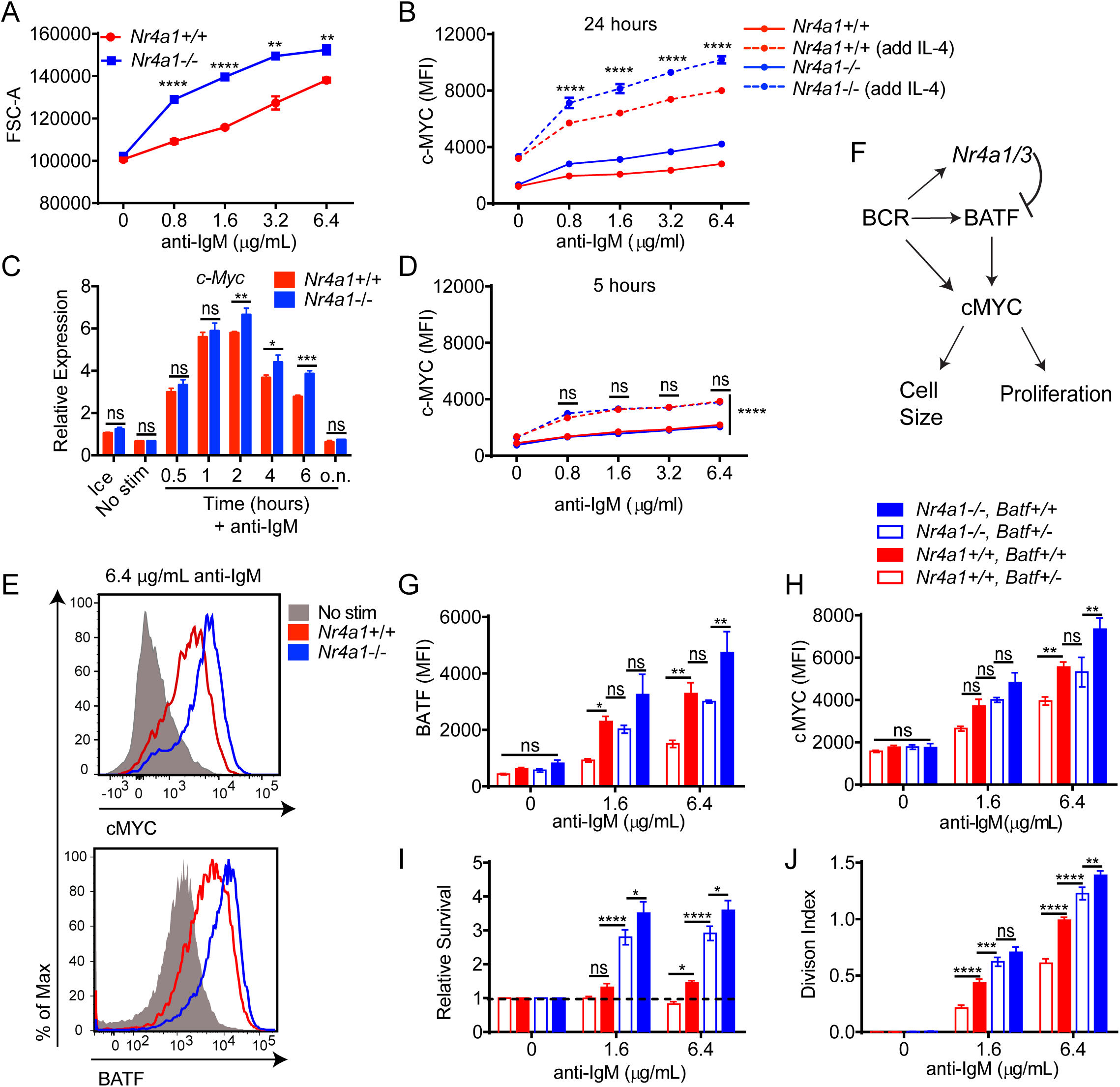
BATF mediates negative regulation of cMYC expression and proliferation by NUR77/*Nr4a1*. **A**. Lymphocytes were harvested from *Nr4a1*+/+ and *Nr4a1*-/- mice, and stimulated with the given dose of anti-IgM for 72 h. Graph depicts forward scatter (FSC) of total live B cells as determined by flow cytometry. **B**. Lymphocytes harvested from *Nr4a1*+/+ and *Nr4a1-/-* mice were stimulated with the given doses of anti-IgM+/- 10ng/ml IL-4 for 24 h. Graph depicts MFI of intracellular c-MYC protein expression in B cells as determined via flow cytometry. **C**. Purified, stimulated B cell samples described in 4A-C and 7G, H were subjected to qPCR to determine relative expression of *cMyc* transcript. **D**. Experiment performed as in (B) except lymphocytes were stimulated for only 6 h. Graph depicts MFI of intracellular c-MYC protein expression in B cells as determined via flow cytometry. **E**. Representative histograms depict intracellular c-MYC and BATF protein expression in gated LN B cells from *Nr4a1*+/+ and *Nr4a1-/-* mice after 24 h stimulation with 6.4 μg/mL anti-IgM. Samples are overlayed onto unstimulated B cells (gray shaded histograms) for reference. **F**. Working model: BCR stimulation drives expression of PRGs including *Nr4a1/3, Batf*, and *cMyc*. c-MYC promotes a broad gene expression program that is required for cell growth and proliferation. Since BATF has been shown to act as a positive regulator of c-MYC, we propose that *Nr4a1/3* gene products restrain BATF and thereby indirectly function as negative regulators of c-MYC and B cell expansion. **G-J**. Lymphocytes harvested from *Nr4a1+/+* and *Nr4a1-/-* mice expressing either 1 or 2 copies of *Batf* (genotypes as shown in legend) were each mixed in a 1:1 ratio with CD45.1+ lymphocytes, +/- CTV loading, and co-cultured in the presence of indicated doses of anti-IgM for either 24 h (G, H) or 72 h (I, J). **G, H**. Graphs depict MFI of either intra-cellular BATF (G) or c-MYC (H) protein expression in CD45.2 B cells. **I, J**. Cells were stained to detect CD45.1, CD45.2, B220 and CTV via flow cytometry. **I**. Shown is the ratio of each CD45.2+ B cells of each genotype relative to co-cultured CD45.1+ WT B cells, normalized to the unstimulated condition. **J**. Graph depicts division index for each genotype. Data in this figure depict N=3 biological replicates for all panels. Mean +/- SEM displayed for all graphs. Statistical significance was assessed with student’s t-test with Holm-Sidak (A, C); two-way ANOVA with Tukey’s (B, D, G-J). *p<0.05, **p<0.01, ***p<0.001, ****p<0.0001

By contrast to c-MYC, the AP-1 family member BATF is the most robustly upregulated transcription factor in *Nr4a1*-deficient B cells at early time points (2 hrs) after Ag stimulation, and is over-induced in B cells lacking all NR4A function (**Figs 8E, 7F, O-R, S7A-C**). We have identified consensus NR4A-binding motifs in regions of open chromatin (OCR; i.e. putative cis-regulatory elements) 6kB upstream and 20kB downstream of *Batf* in mature follicular B cells (ImmGen ATAC-seq data), suggesting that *Batf* is likely to be a direct transcriptional target of NR4A family. Since BATF can cooperate with IRF4 to induce *cMyc* expression^64^, and *Batf*-deficient B cells exhibit defective clonal expansion^65^, we hypothesized that NUR77 regulates c-MYC in part via modulation of BATF (see model, **Fig 8F**). To test this hypothesis, we sought to rescue BATF over-induction in NUR77-deficient B cells by generating *Nr4a1-/- Batf+/-* mice. We found, as expected, that hemizygosity for *Batf* reduced expression by approximately 50% and resulted in a near-normalization of BATF protein expression in *Nr4a1*-deficient B cells after BCR stimulation (**Fig 8G, S8D**). As predicted by our model (**Fig 8F**), c-MYC over-induction was concommitantly rescued (**Fig 8H**). Across genotypes, we observed a correlation between BATF and c-MYC expression, and B cell proliferation (**Fig 8G-J**). Importantly, in support of our working model, dose reduction of *Batf* partially rescued excess proliferation (but not enhanced survival) of *Nr4a1-/-* B cells (**Fig 8I, J, S8E, F**).

### NUR77/*Nr4a1* restrains expression of target genes that facilitate recruitment of T cell help by B cells

Although we identified a role for the NR4A family in selectively regulating B cell expansion in response to Ag stimulation (signal 1) without co-stimulation (signal 2) (**Figs 2, 3, 5, 6**), the most differentially induced genes in *Nr4a1-/-* B cells were *Ccl3* and *Ccl4*, which encode chemokines that serve to recruit T cell help via CCR5 (**Fig 7C-E, G-N**)^51,53,57^. In addition, RNAseq analysis identified additional putative NUR77 target genes that are well-recognized to facilitate engagement of T cell help for B cells, including *Cd86* and *Icam1* (**Fig 7C, Supplementary Data 1)**. CD86 is a ligand for CD28, while ICAM1 plays a role in supporting B cell-T cell conjugates via LFA1 engagement, and its expression together with ICAM2 is important for efficient expansion of B cells and seeding of the germinal center in response to T-dependent immunization^2,49^. We validated these candidate target genes by assessing protein upregulation in response to anti-IgM stimulation. CD86 is over-induced in the absence of *Nr4a1* but not *Nr4a3*, and this is recapitulated in cDKO B cells in a cell-intrinsic manner (**Fig S9A, B**). Similarly, ICAM1 is over-induced in Ag-stimulated *Nr4a1-/-* B cells (**Fig S9C, D**). Collectively, these data suggested to us that the NR4A family may play a role in T-dependent immune responses, particularly in situations where competition for limiting quantities of T cell help is important.

### NUR77/Nr4a1 restrains B cell access to limiting amounts of T cell help under competitive conditions *in vivo*

Although we showed that NUR77/*Nr4a1* expression did not influence B cell responses to co-stimulatory signals *in vitro* and *in vivo* (**Figs 3, 6**), we reasoned that access to T cell help (signal 2) in these assays was not limiting. Since we identified over-induction of target genes that play important roles in the recruitment and engagement of T cell help in Ag stimulated *Nr4a1-/-* B cells (*Ccl3, Ccl4, Cd86, Icam1*) (**Fig 7, S9A-D**), we hypothesized that *Nr4a1*-deficient B cells might exhibit a competitive advantage in the context of a limiting supply of T cell help.

To test this hypothesis, we devised an experimental strategy in which we could modulate the amount of T cell help available to Ag specific B cells *in vivo*. To do so we again undertook adoptive transfer of Ag-specific *Nr4a1+/+* or *Nr4a1-/-* B cells harboring the B1-8i HC Tg. To ensure dependence upon T cell help, we transferred congenically marked purified B cells into hosts one day prior to immunization with the T-dependent immunogen NP-OVA in order to allow proper localization of donor B cells prior to Ag encounter (**Fig 9A, 6E, F**). By co-transferring varying numbers of OVA-specific OTII splenocytes in conjunction with donor B cells into either WT or *CD40L-/-* hosts, we could manipulate the supply of Ag-specific T cell help (**Fig 9B**). As expected, we observed a profound defect in Ag-specific B cell proliferation in *CD40L-/-* hosts that could be rescued with adoptive transfer of Ag-specific T cells (**Fig 9C**).

**Figure 9.**
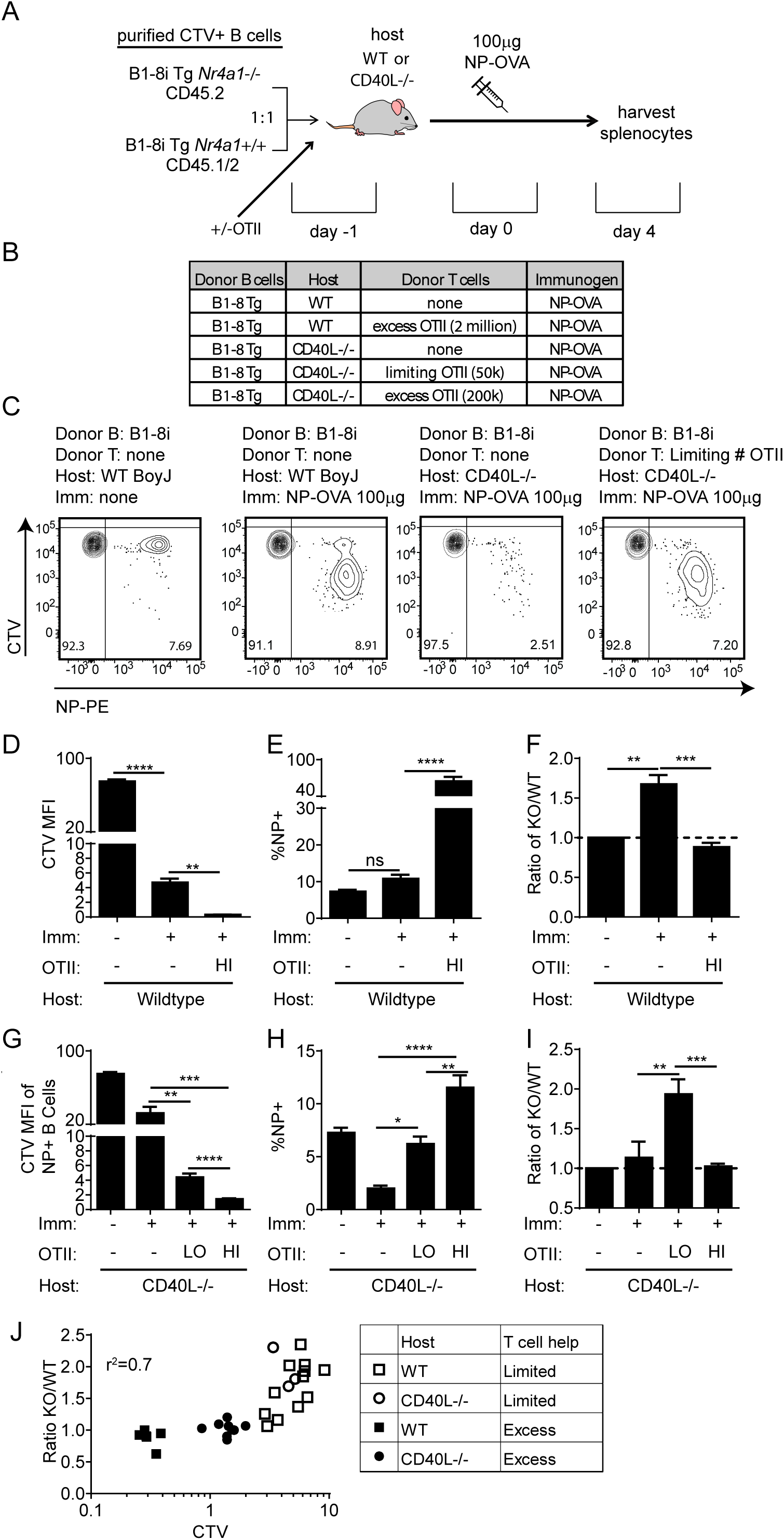
NUR77/Nr4a1 restrains B cell competition for T cell help under limiting conditions. **A, B**. Schematic of adoptive transfer experimental design. B cells were purified from splenocytes harvested from *Nr4a1*+/+ B1-8 Tg CD45.1/2+ and *Nr4a1*-/- B1-8 Tg CD45.2+ mice via bench-top negative selection, mixed 1:1, and loaded with CTV. 2×10^6^-3×10^6^ B cells were then adoptively transferred +/- OTII splenocytes into either WT or *CD40L-/-* hosts. Host mice were then immunized IP one day later with 100 µg NP-OVA/alum, followed by spleen harvest on d4. **C**. Representative flow plots gated on donor B cells depict CTV dilution among NP-binding B cell under varied conditions. **D-F**. Adoptive transfers into WT hosts were performed as described above (A, B). Mice which excess T cell help (OTII: “HI”) received donor B cells co-transferred with 2×10^6^ OTII splenocytes Graphs depict CTV dilution among donor NP-binding B cells (D), expansion of NP-binding donor B cells (E), and ratio of *Nr4a1*-/- relative to *Nr4a1*+/+ donor NP-binding B cells normalized to the ratio in unimmunized hosts (F). N=5 unimmunized recipients, N=13 hosts that received only donor B cells, and N=6 hosts that received donor B cells mixed with excess OTII. **G-I**. Adoptive transfers into *CD40L-/-* hosts were performed as described above (A, B). Mice which received limited T cell help (OTII: “LO”) received donor B cells co-transferred co-transferred with 5×10^4^ splenocytes harvested from OTII mice, and mice which received excess T cell help (OTII: “HI”) received donor B cells co-transferred co-transferred with 2×10^5^ splenocytes. Graphs depict CTV dilution among donor NP-binding B cells (G), expansion of NP-binding donor B cells (H), and ratio of *Nr4a1*-/- relative to *Nr4a1*+/+ donor NP-binding B cells normalized to the ratio in unimmunized hosts (I). N=5 unimmunized recipitents, N= 7 hosts that received B1-8 donor B cells only, N=3 hosts that received limited T cell help, and N=9 hosts that received excess OTII T cells. **J**. Graph depicts correlation between ratio of *Nr4a1*-/- relative to *Nr4a1*+/+ donor NP-binding B cells (as plotted in F, I above) and CTV dilution of NP-binding donor B cells in individual recipients (as a proxy measure of T cell help). Mean +/- SEM displayed for all graphs. Statistical significance was assessed with one-way ANOVA with Tukey’s (D-G, I) or Sidak (H), and Pearson correlation coefficient (J). *p<0.05, **p<0.01, ***p<0.001, ****p<0.0001

Although we observed robust vital dye dilution and expansion of adoptively transferred Ag-specific B cells in WT hosts after immunization with NP-OVA, this response increased dramatically with provision of excess T cell help, suggesting that endogenous OVA-specific T cells are limiting (**Fig 9D, E**). Indeed, we observe a striking competitive advantage for *Nr4a1-/-* NP-specific donor B cells in immunized WT hosts that is completely lost with provision of excess T cell help (**Fig 9F**). We next sought to reproduce this finding in *CD40L-/-* mice where endogenous T cell help is defective (**Fig 9C**). Upon transfer of either no, limiting, or excess numbers of OTII splenocytes into *CD40L-/-* hosts, the amplitude of donor B cell expansion scaled with the amount of Ag-specific T cell help (**Fig 9G, H**). Again, we observed a clear competitive advantage for Ag specific *Nr4a1*-/- B cells specifically under conditions where T cell help is limiting, that is in turn completely lost with provision of excess T cell help (**Fig 9I**).

We next took advantage of heterogeneity in the amplitude of B cell responses in individual host mice to look for a correlation between the supply of T cell help and any competitive advantage enjoyed by *Nr4a1-/-* B cells. To do so, we used vital dye dilution of Ag-specific donor B cells in individual hosts as a proxy measure of the supply of T cell help. Indeed, we identify a clear inverse correlation (r^2^ = 0.7) between the supply of T cell help and competitive fitness of Ag-specific *Nr4a1-/-* donor B cells (**Fig 9J, S9E**). These data suggest that NUR77 mediates a negative feedback loop downstream of Ag stimulation, and may play a role not only in T-independent responses, but also in T-dependent immune responses, specifically in competitive settings where T cell help is limiting.

## DISCUSSION

Ag-activated B cells that fail to recruit cognate T cell help within a narrow time window either undergo apoptosis, become anergic, or revert to a “naïve-like” state, depending on the strength and duration of Ag stimulation^4,5^. Our data suggest that induction of *NR4A* gene expression represents a novel molecular strategy by which B cells acutely activated only by Ag are normally restrained, and helps to enforce their dependence upon co-stimulation. Indeed, we have previously identified upregulation of NUR77-EGFP expression in self-reactive B cells in response to chronic BCR engagement, and isolated a specific role for NUR77 in restricting the survival of such cells, particular in settings where the B cell survival factor BAFF is limiting^25,27-29^. It has been proposed that certain features of anergic B cells, including reduced half-life, may be understood in part as a “special case” of the acute B cell response to signal 1 in the absence of signal 2^66^. The NR4A family may supply one example of a shared molecular link between these two phenomena.

Importantly, there is a physiologic temporal delay between initial Ag encounter and acquisition of co-stimulatory input in the form of T cell help; co-stimulatory signals must be received within a limited span of time in order to divert Ag-stimulated B cells from apoptosis/anergy and facilitate their recruitment into humoral immune responses^4,5^. Hodgkin and colleagues have long postulated the existence of a ‘death timer’ in activated B cells^61^. Recent work from Akkaya, Pierce and colleagues described the gradual accumulation of intracellular calcium in Ag-activated B cells that is accompanied by progressive development of irreversible mitochondrial dysfunction and gradual loss of glycolytic capacity^5^. These processes may serve as a molecular timer that marks the “point of no return” after which B cells can no longer be rescued from apoptosis by co-stimulation^5^. Consistent with previous observations, this time window scales with both the duration and strength of Ag stimulation^4,5^.

We show that *Nr4a1* and *Nr4a3* cooperatively promote Ag-induced apoptosis, and this is rescued with provision of co-stimulation (**Figs 2, 3, 5, S5**). We hypothesize that the balance between NR4A-induced apoptosis and pro-survival signals generated by BCR ligation may help to modulate the time window within which T cell help must be engaged to avoid a terminal fate. Indeed, *Nr4a1-/-* B cells exhibit an advantage relative to WT B cells when provision of co-stimulatory signals is delayed, suggesting precisely such a role (**Fig S3G**). NUR77 has been shown to trigger apoptosis by inducing a conformational change that exposes the pro-apoptotic BH3-only domain of BCL-2 and related family members^11,12^. It will be important to determine whether and how this interaction is modulated by BCR and co-stimulatory signals, and whether this intersects with the progressive metabolic dysfunction described by Akkaya and colleagues, or instead represents a parallel, independent mechanism to restrain self-reactive B cells from mounting inappropriate responses to endogenous Ags^5^.

Direct activation of apoptosis may not be the only mechanism by which the NR4A family promotes antigen-induced cell death; since the initial increase in the glycolytic capacity of Ag-activated B cells depends on cMYC induction, we propose that the NR4A family may contribute to a gradual loss in glycolytic capacity and metabolic dysfunction over time by suppressing cMYC induction (**Fig 8, S8**)^62^. Of note, in contrast to naïve B cells that encounter Ag, anergic B cells remain metabolically quiescent in the face of Ag stimulation^62^. We therefore speculate that NR4A expression in self-reactive B cells, as described in our prior studies, might also contribute to this phenomenon by repressing inducible cMYC expression^25,28^.

Although we show that the competitive advantage exhibited by *Nr4a1-/-* B cells is lost with provision of co-stimulation (**Fig 3**), endogenous NUR77 induction exhibits comparable kinetics either with or without addition of signal 2 (**Fig 1F, S3A-C**). This raises the question of how co-stimulation circumvents inhibitory effects of the NR4A family. One possible explanation is that co-stimulation provides robust pro-survival and proliferative signals (mediated in part by a profound increased in cMYC protein levels) that overwhelm the inhibitory tone supplied by the NR4A family (**Fig 8B**). Another potential mechanism (that is not mutually exclusive) is that post-translational modifications of NR4A family members triggered by Ag and/or co-stimulatory ligands could regulate its subcellular localization or protein-binding interactions to influence downstream effector functions such as apoptosis or target gene transcription^6,67-70^. Importantly, we still observe over-induction of c-MYC in *Nr4a1-/-* B cells even in the context of co-stimulation, suggesting that repression of at least some target genes by NUR77 is not altered by co-stimulatory signals (**Fig 8B**).

The initial transcriptional program elicited by Ag encounter in B cells serves to drive cell cycle entry, metabolic reprogramming, and facilitates recruitment of T cell help^2^. Indeed, even a “single round” of BCR stimulation is sufficient to trigger these events^3^. Strikingly - similar to other mitogens - BCR-induced PRG expression is transient. It has long been appreciated that chemical inhibition of new protein synthesis results in ‘super-induction’ and sustained transcript expression of mitogen-induced PRGs^71,72^. This implies the existence of one or more newly translated negative feedback regulators downstream of mitogenic stimuli that inhibit new transcript synthesis and reduce transcript stability^73^. Indeed, several such factors have been described in other contexts, but the identity of such negative feedback regulators in B cells remains to be defined^48,73^. We propose that the NR4A family represent excellent candidates to contribute to this phenomenon and in fact do so for a small subset of target genes including *Ccl3, Ccl4*, and *Batf* (**Figs 7G, H, O**). Elucidation of other pathways that account for ‘super-induction’ in B cells will be very interesting to explore.

In this manuscript, we have identified and validated several novel NR4A targets in Ag-activated B cells. Collectively, the subset of B cell PRGs that are negatively regulated by the NR4A family play important roles in orchestrating humoral immune responses^49-56,65^. Several target genes play a specific role in T cell–B cell interactions. Most notably, secretion of the chemokines CCL3 and CCL4 (MIP-1 alpha and beta) by B cells can induce both activated conventional and regulatory T cell migration via CCR5^51,53,57,74^. CD86 has a very well-established role in T cell co-stimulation, and suppression of CD86 upregulation represents a key checkpoint in anergic B cells^75,76^, breach of which is sufficient to break tolerance^77^. ICAM1 and 2 facilitate T-B conjugate formation during T-dependent immune responses and are critical for efficient B cell expansion in this context^49^.

Other critical mediators of the humoral immune response are also regulated by the NR4A family. B cell-derived VEGF-alpha can drive lymphangiogenesis, expansion of LNs and the development of high endothelial venules^50^. CD69 is a well-recognized ‘activation’ marker on lymphocytes, and functions to regulate retention of lymphocytes in secondary lymphoid organs via negative regulation of S1P1^52^. Although the zinc finger TFs EGR2 and EGR3 have tolerogenic and regulatory functions in the setting of chronic Ag stimulation, like EGR1, they also play important roles in response to acute Ag stimulation^54,78^. Indeed, both TCR and BCR-induced proliferation are defective in *Egr2/3* cDKO lymphocytes, and this has been attributed in part to direct regulation of *c-Myc*^79^. BATF heterodimerizes with JUN to bind cooperatively with IRF4 at so-called AP-1-IRF4 composite elements (AICE)^80-82^, and as a result, BATF plays pleiotropic roles as a transcriptional regulator in immune cells; it is required for Th17 and Tfh differentiation, and is essential for CSR in B cells^38,55,56^. By virtue of cooperative binding with IRF4 (and regulation of *c-Myc* along with other targets), BATF may serve as a key regulatory node that helps to translate signal strength into B cell fate decisions^83,84^. Indeed, its deletion has profound consequences for early and late B cell responses to T-dependent immunogens^65^. We therefore speculate that repression of *Batf* expression (along with *Egr2/3*) may in turn mediate broad modulation of gene expression programs and B cell fate by NUR77 and its family members.

The literature provides compelling evidence for functional redundancy between NR4A family members in immune cells^13-16,21^. Perhaps the most striking example of this comes from genetic studies showing that conditional deletion of multiple *NR4A* genes in thymocytes results in complete loss of Tregs and development of a severe *scurfy*-like autoimmune disease, while mice deficient for individual family members remain healthy ^15^. We reproduce this finding with our novel germline *Nr4a3-/-* model and go on to take advantage of conditional *Nr4a1*^*fl/fl*^ mice in order to delete both family members in B cells (cDKO model) while avoiding systemic consequences of global *Nr4a* gene deletion. We detect extremely low *Nr4a2* transcript abundance even in Ag-stimulated B cells and minimal over-induction in our cDKO model (**Fig 1D, 4I**). We therefore conclude that virtually all NR4A function is lost in cDKO B cells. We show that although *Nr4a3-/-* B cells exhibit little to no over-induction of NUR77 target genes, cDKO B cells lacking both *Nr4a1* and *Nr4a3* show synergistic effects of deleting both family members, suggesting that *Nr4a1* and *Nr4a3* indeed play partially redundant roles in B cells (**Fig 7, S7**). Similarly, we observe *NR4A* gene dose-dependent increases in survival and proliferation of Ag-stimulated B cells (**Fig 2E, 5**). Interestingly, *Nr4a3-/-* B cells exhibit enhanced proliferation relative to WT B cells despite minimal change in c-MYC expression suggesting that NOR1/*Nr4a3* may also have unique and non-redundant transcriptional targets in B cells. Although we cannot formally exclude a B cell-extrinsic contribution of *Nr4a3* deletion in cDKO B cell phenotypes, we do reproduce target gene over-induction and *in vitro* functional phenotypes in a 1:1 mixed chimera. Moreover, unperturbed candidate target gene expression in *Nr4a3-/-* B cells also suggests our observations in cDKO mice reflect B cell-intrinsic effects of *Nr4a1* and *Nr4a3*.

Although provision of co-stimulation bypassed inhibitory functions of the NR4A family *in vitro* and *in vivo*, several key target genes that are over-induced in the absence of NR4A expression (*Ccl3, Ccl4, Cd86, Icam1*) play well-established roles specifically in the recruitment and engagement of T cell help^2,49,51,53,57^. Indeed, we showed that *Nr4a1-/-* B cells exhibit a competitive advantage specifically in the setting of limiting (but not excess) T cell help *in vivo* (**Fig 9**). Since NR4A expression scales with the intensity of Ag stimulation, we predict that the NR4A family may disproportionately restrain the most strongly Ag-activated B cell clones in a polyclonal repertoire. This would allow weaklys-stimulated lower affinity clones to participate in a humoral immune response, and thereby limit immunodominance of high affinity clones. A specific mechanism to limit immunodominance is especially critical in the context of early humoral immune responses in order to preserve clonal diversity as germinal centers form; excessive immunodominance of a single high affinity clone early during a humoral immune response severely compromises the efficiency of affinity maturation^85^. Indeed, although B cell clones compete for clonal dominance and limited resources (T cell help) during the primary immune response^86^, low affinity B cell clones do enter (and persist in) the germinal center^87,88^, suggesting that some physiological mechanism must exist to restrain high affinity B cell clones from completely monopolizing T cell help^87^. We propose that the NR4A family may serve this role by mediating a negative feedback loop downstream of the BCR. Future work will be important to further test this hypothesis directly.

Although no endogenous ligand has been identified for the NR4A family, several small molecule agonist and antagonist ligands for NUR77/*Nr4a1* have been described^89-92^. Therefore, targeting specific members of the NR4A family may be a viable therapeutic approach for a range of immune-mediated diseases. Indeed, agonist ligands for the NR4A family are being actively developed for clinical use in treatment of hematologic malignancies^93^ and deletion of NR4A family members may augment CAR T cell clearance of tumor cells^16,17^. One could envision targeting B cell-mediated autoimmune disease with a selective NUR77 agonist. Conversely, an antagonist compound could boost T-independent B cell responses and serve as a novel ‘universal adjuvant’. The ability to selectively target individual NR4A family members may make these types of approaches feasible in the future.

In summary, here we identify a novel negative feedback loop downstream of Ag stimulation that is mediated by the NR4A family and serves to restrains B cells that receive signal 1 in the absence of signal 2. We postulate that this functions as a B cell tolerance mechanism that renders B cells dependent upon receipt of co-stimulatory input within a defined time window. Unexpectedly, we also show that *Nr4a1-/-* B cells exhibit a competitive advantage under conditions of limiting T cell help. We propose that this mechanism serves to restrain high affinity B cell clones from monopolizing limiting quantities of T cell help in order to limit immunodominance and preserve clonal diversity. Our work thus reveals a new molecular mechanism by which self- and foreign-reactive B cells are regulated and may be therapeutically manipulated.

## Supporting information

Supplemental Data 1_RNAseq

## ACKNOWLEDGEMENTS

We thank Al Roque for help with mouse husbandry. We thank Arthur Weiss, Mark Ansel, Jeroen Roose, and Christopher Allen for critical scientific feedback. We thank our funders: NIAID 5T32AI007334-28 (CT), HHMI Medical Research Fellows program (JH), NIAMS R01AR069520 (JZ), and Rheumatology Research Foundation (JZ). A.M. holds a Career Award for Medical Scientists from the Burroughs Wellcome Fund, is an investigator at the Chan Zuckerberg Biohub and a member of the Parker Institute for Cancer Immunotherapy (PICI) and has received funding from the Innovative Genomics Institute (IGI).

## DECLARATION OF INTERESTS

The authors declare competing financial interests: A.M. is a co-founder of Arsenal Biosciences and Spotlight Therapeutics. A.M. serves as on the scientific advisory board of PACT Pharma, is an advisor to Trizell, and was a former advisor to Juno Therapeutics. The Marson Laboratory has received sponsored research support from Juno Therapeutics, Epinomics, Sanofi and a gift from Gilead. J.Z. serves as a scientific consultant for Walking Fish Therapeutics.

## Notes

https://www.ncbi.nlm.nih.gov/geo/query/acc.cgi?acc=GSE146747

