## Supplemental Figure legends, Table, Figures for "“A negative feedback loop mediated by the NR4A family of nuclear hormone receptors restrains expansion of B cells that receive signal one in the absence of signal two”"

**Supplemental Figure 1. A.** Splenocytes from *Nr4a1*+/+ and *Nr4a1-/-* mice were surface stained to identify CD23+ B cells and loaded with Indo-1 dye. Samples were collected on a flow cytometer for 20 seconds to assess basal calcium flux, and then for a total of 3m following stimulation with either 5 µg/mL anti-IgM or a 1:200 dilution of polyclonal anti-IgD. Ratiometric assessment of intracellular calcium is depicted for CD23+ B cells. **B, C.** Splenocytes from *Nr4a1*+/+ and *Nr4a1-/-* mice were stimulated with the given doses of anti-IgM for 5 min, fixed, permeabilized, and then stained intracellularly for either phospho-Erk (B) or phospho-S6 (C) followed by surface staining to identify B cells. Histograms depict overlays of CD23+ B220+ B cells of each genotype across doses. **D.** Splenocytes from congenically marked *Nr4a1*+/+ and *Nr4a1-/-* mice were mixed 1:1, and then either stimulated with 10 µg/mL anti-IgM or incubated with media alone for 4 hours. The stimulus was then washed out, and the cells were rested in media alone for 15 minutes. Finally, the cells were re-stimulated with the given dose of anti-IgD for five minutes and then, fixed and stained as in B, C to detect intra-cellular phosphor-Erk. Data in this figure reflect N=2 biological replicates for all panels.

**Supplemental Figure 2. A.** Lymphocytes from either *Nr4a1*+/+ or *Nr4a1*-/- CD45.2+ mice were mixed in a 1:1 ratio with CD45.1+ lymphocytes, CTV loaded and co-cultured in the presence of indicated doses of anti-IgM +/- 20 ng/mL BAFF for 72 h. Cells were then stained to detect CD45.1, CD45.2, B220, CTV via flow cytometry. Samples correspond to those depicted in Fig 2A, B. Graph depicts total live B cell number in each sample as calculated via flow cytometry in reference to a fixed number of fluorescent counting beads. **B.** Lymphocytes from NUR77-EGFP reporter mice were stimulated with the indicated doses of anti-IgM +/- 20 ng/mL BAFF for 24 h. NUR77-EGFP MFI of live B cells was determined via flow cytometry. **C-F.** Lymphocytes were harvested from CD45.2+ *Nr4a1*+/+ or *Nr4a1*-/- mice, mixed 1:1 with CD45.1+ WT lymphocytes, and co-cultured with the given doses of anti-IgM +/- 20 ng/mL BAFF for either 24 or 48 hours. **C, E.** Graphs depict the % of B cells expressing activated caspase-3 as assessed by intracellular staining via flow cytometry in samples without (C) or with (E) addition of BAFF. **D, F**. Graphs depict the ratio of CD45.2+ *Nr4a1*-/- or *Nr4a1+/*+ B cells relative to

CD45.1+ co-cultured WT B cells normalized to the input ratio in samples without (D) or with (F) addition of BAFF. **G.** Splenocytes were harvested from *Nr4a1*+/+, *Nr4a1*+/- and *Nr4a1*-/- mice, and stimulated with 10 μg/mL anti-IgM for 2 hours. Graph depicts MFI of endogenous NUR77 expression in CD23+ B cells as determined by intracellular staining via flow cytometry. **H.** Either purified B cells obtained by bench-top negative selection or total lymphocytes from CD45.2+ *Nr4a1*-/- and CD45.1+ *Nr4a1+/+* mice were mixed 1:1 and co-cultured with 6.4μg/ml anti-IgM for 72 h. Representative histograms depict CTV dilution in B cells under each condition and are representative of at least 4 biological replicates. **I-J.** Lymphocytes harvested from either *Nr4a1*+/+ or *Nr4a1*-/- CD45.2+ IgHEL Tg mice were mixed 1:1 with CD45.1 WT lymphocytes, CTV loaded, and co-cultured in the presence of indicated doses of anti-IgM +/- 20 ng/mL BAFF for 72 h. Cells were then stained to detect CD45.1, CD45.2, B220, CTV via flow cytometry. **I**. Shown is the ratio of either *Nr4a1*+/+ or *Nr4a1*-/- CD45.2+ IgHEL Tg B cells relative to co-cultured CD45.1+ WT B cells, normalized to the input ratio. **J.** Graph depicts division index of CD45.2+ B cells for each genotype. **K, L**. Samples correspond to those described in Fig 2H, cultured in the presence of 20ng/ml BAFF. **K.** Displayed is the ratio of cre+ or cre+ *Nr4a1* fl/fl CD45.2+ B cells relative to co-cultured CD45.1+ WT B cells, normalized to the input ratio. **L.** Graph depicts division index of each genotype. **M, N.** Lymphocytes from CD45.2+ mice with either mb1-cre only, conditional, or germline deletion of *Nr4a1* (mb1-cre *Nr4a1 fl/fl* or *Nr4a1-/-*) were mixed 1:1 with CD45.1+ lymphocytes, CTV loaded and co-cultured in the presence of indicated doses of anti-IgM for 72 h. Cells were then stained to detect CD45.1, CD45.2, B220 and CTV via flow cytometry. **M.** Graph depicts total B cell numbers of each genotype as determined by flow cytometry using fixed number of labeled beads as a reference. **N.** Graph depicts division for each genotype. Data in this figure depict N=3 biological replicates for all panels except (H) as noted. Mean +/- SEM displayed for all graphs. Statistical significance was assessed with student's t-test with Holm-Sidak (A-F, I-L); two-way ANOVA with Tukey's (M, N). *p<0.05, **p<0.01, ***p<0.001, ****p<0.0001

**Supplemental figure 3. A-C.** Lymphocytes from WT mice were stimulated with the given doses of the indicated stimulus for the given length of time. Graphs depict endogenous NUR77 expression in B cells as determined via intracellular flow staining. **D-F.** Lymphocytes from CD45.1+ *Nr4a1+/+* and CD45.2+ *Nr4a1-/-* mice were mixed 1:1, loaded with CTV, and co-cultured for 72 h with the given stimulus. Cells were then stained to detect CD45.1, CD45.2, B220, CTV via flow cytometry. **D.** Representative histograms depict CTV dilution of *Nr4a1+/+* and *Nr4a1*-/- B cells stimulated with: 10 µg/mL LPS, 2 µM CPG, or 10 µg/mL anti-IgM +/- either 1 µg/mL anti-CD40 or 10 ng/mL IL-4. **E.** Graph depict % of B cells that divided at least once as determined by CTV dilution. **F.** Graph depicts ratio of CD45.2+ *Nr4a1-/-* B cells relative to co-cultured CD45.1+ *Nr4a1+/+* B cells, normalized to the input ratio. **G.** Mixed co-cultures were generated as described above in (D-F) in the presence of low dose anti-IgM 1 µg/mL. Co-stimulation with 1 µg/mL anti-CD40 was added at the indicated timepoint, and cells were harvested for analysis after a total of 72 h in culture each condition. Shown are representative histograms depicting CTV dilution of live B cells of each co-cultured genotype. Histograms are representative of 4 biological replicates. Graphs in this figure depict N=3 biological replicates for all panels. Mean +/- SEM displayed for all graphs. Statistical significance was assessed with student's t-test with Holm-Sidak (E); one-way ANOVA with Dunnett's (F). *p<0.05, **p<0.01, ***p<0.001, ****p<0.0001

**Supplemental Figure 4. A.** Purified, stimulated B cell samples described in Fig 4A-C were subjected to qPCR to determine relative expression of *GFP* transcript. **B.** Lymphocytes from *Nr4a1+/+* and *Nr4a1-/-* mice which expressed the NUR77-EGFP reporter were stimulated with the given doses of anti-IgM for 24 h. Graph depicts GFP MFI in B cells as determined by flow cytometry. **C, D.** B cells from *Nr4a1*+/+, *Nr4a1-/-*, and *Nr4a3-/-* mice were purified by bench-top negative selection from pooled splenocytes, and were stimulated with 10 µg/mL anti-IgM for the indicated times. qPCR was performed to determine relative expression of *Nr4a3* (C) and *Nr4a1* (D) transcripts. **E.** Thymocytes from WT, *Nr4a1*-/- and *Nr4a3*-/- mice were incubated +/- PMA and ionomycin for 2 h. Subsequently whole cell lysates were blotted with Ab to detect NUR77, NOR1, c-MYC, and GAPDH. Red arrow indicates presence of low abundance truncated NOR1 protein in *Nr4a3-/-* mice. **F, H, I.** Splenocytes from WT, *Nr4a1-/-* and *Nr4a3-/-* mice were stained to detect T cell subsets on the basis of CD64L and CD44 expression. **F.** Representative plots of CD4+ (top row) and CD8+ (bottom row) T cells depict gating to identify naïve and memory populations. CM=central memory. EM=effector memory. **G.** Thymocytes and splenocytes from WT, *Nr4a1-/-* and *Nr4a3-/-* mice were permeabilized and stained to detect intra-cellular Foxp3 to identify Treg. Graph depicts % Tregs in CD4+ gates in thymus and spleen. **H, I.** Graphs depict quantification of T cell subsets as gated in (F). **J, K.** Splenocytes from WT, *Nr4a1-/-* and *Nr4a3-/-* mice were stained to detect B cell subsets on the basis of CD21, CD23, and CD93 expression. **J.** Representative plots of B220+ cells (top row) to identify marginal zone (MZ) population. Bottom row depicts T1, T2/3, and Fo gates drawn on B220+ cells after exclusion of MZ. **K.** Graph depicts quantification of B cell subsets as gated in (J). **L, M.** Chimeras were generated as described in Fig 4H. Graphs depict quantification of T cell subsets as gated in (F). N=5 chimeras / genotype. **N-O.** Irradiated CD45.1+ recipients were reconstituted with 1:1 bone marrow mix from donor A (CD45.1/2 WT mice) and donor B (either CD45.2+ mb1-cre control or CD45.2+ cDKO mice). Chimeras were sacrificed 10-12 weeks after reconstitution. Relative reconstitution of B cell developmental subsets in the bone marrow and spleen was determined via flow cytometry. N=5 for chimeras / mix. Graphs in this figure depict N=3 biological replicates for all panels except (L-O) as noted above. Mean +/- SEM displayed for all graphs. Statistical significance was assessed with student's t-test with Holm-Sidak (A, B); two-way ANOVA with Tukey's (C, D, G-K); student's t-test without correction (L, M). *p<0.05, **p<0.01, ***p<0.001, ****p<0.0001

**Supplemental figure 5. A, B.** Samples correspond to those described in Fig 5B, C, cultured in the presence of 20 ng/ml BAFF. **A.** Shown is the ratio of CD45.2+ WT, *Nr4a1-/-* or *Nr4a3-/-* B cells relative to co-cultured CD45.1+ WT B cells, normalized to the unstimulated condition. **B.** Graph depicts division index for each genotype. **C, D.** Samples correspond to those described in Fig 5F, G, cultured in the presence of 20 ng/ml BAFF. **C.** Shown is the ratio of each CD45.2+ B cell genotype relative to co-cultured CD45.1+ WT B cells, normalized to the unstimulated condition. **D.** Graph depicts division index for each genotype. **E-J.** Competitive bone marrow chimeras were generated as described in Fig S4N, O. 10-12 weeks after reconstitution, lymphocytes were harvested from chimeras, CTV loaded and cultured with given stimuli (anti-IgM doses +/- 20 ng/ml BAFF +/- 10 ng/ml IL-4). N=5 chimeras of each genotype were analyzed. **E, G, I.** Graphs depict ratio of CD45.2+ cre+ or cDKO B cells relative to CD45.1/2 WT B cells from each chimera, normalized to unstimulated condition. **F, H, J.** Graphs depict division index for CD45.2+ cre+ or cDKO B cells from each chimera. Graphs in this figure depict N=3 biological replicates for panels (A-D) except (E-J) as noted above. Mean +/- SEM displayed for all graphs. Statistical significance was assessed with two-way ANOVA with Tukey's (A-D); student's t-test (E-J). *p<0.05, **p<0.01, ***p<0.001, ****p<0.0001

**Supplemental Figure 6. A.** *Nr4a1*+/+ and *Nr4a1*-/- mice were immunized IP with 100 μg NP-LPS (B) and anti-NP IgM titers were determined via ELISA at serial time points. Samples correspond to those in Fig 6B. **B.** Adoptive transfer and immunization performed as described in Fig 6E-G. Shown is the ratio of recovered donor *Nr4a1*-/- B1-8 Tg CD45.2+ B cells relative to donor *Nr4a1*+/+ B1-8 Tg CD45.1/2+ B cells that fall either into the NP-binding or non-NP binding gates as shown in Fig 6F. Data depicted correspond to Fig 6G. **C.** Splenocytes harvested from *Nr4a1*+/+ B1-8 Tg CD45.1/2+ and *Nr4a1*-/- B1-8 Tg CD45.2+ cells were mixed 1:1 and loaded with CTV. $5 \times 10^6$ cells were adoptively transferred into WT hosts. Hosts were immunized IP on the same day with 10 μg/mL NP-KLH/alum and an equal number of control hosts were left unimmunized, followed by spleen harvest on d3. Shown is the ratio of *Nr4a1*-/- to *Nr4a1+/+* donor B cells among the NP-binding population, normalized to input ratio. Data in A depicts N=4-5 mice /genotype, and data in B, C depicts N=3-4 recipient mice per condition. Mean +/- SEM displayed for all graphs. Statistical significance was assessed with student's t-test. *p<0.05, **p<0.01

**Supplemental Figure 7. A.** Purified, stimulated B cell samples described in 4A-C and 7G, H were subjected to qPCR to determine relative expression of *Batf* transcript. **B.** Lymphocytes harvested from *Nr4a1+/+* and *Nr4a1-/-* mice were stimulated with the given doses of anti-IgM for 24 hours. Graph depicts MFI of intracellular BATF protein expression in B cells as determined via flow cytometry. **C.** Competitive bone marrow chimeras were generated as described in Fig S4N, O. After reconstitution, lymphocytes harvested from the chimeras were stimulated with the given doses of anti-IgM for 24 h. Graph depicts MFI of intracellular BATF protein expression in CD45.2+ Cre-only or cDKO B cells as determined via flow cytometry. **D.** RNAseq performed as described in Fig 7A. Displayed is the FPKM of *Cd69*. **E.** Lymphocytes harvested from CD45.1+ *Nr4a1+/+* and CD45.2+ *Nr4a1-/-* mice were mixed 1:1 and co-cultured with the given doses of anti-IgM for 24 hours. Graph depicts MFI of CD69 surface expression on each genotype of B cells as determined via flow cytometry. **F.** Purified, stimulated B cell samples described in Fig 7K, L were subjected to qPCR to determine relative expression of *Cd69* transcript. **G**. Experiment performed as in Fig 7P. Graph depicts MFI of CD69 surface expression on B cells as determined via flow cytometry. **H.** Purified, stimulated B cell samples described in Fig 7M, N were subjected to qPCR to determine relative expression of *Cd69* transcript. **I**. Experiment performed as in Fig 7R. Graph depicts MFI of CD69 surface expression on B cells as determined via flow cytometry. **J.** Experiment performed as in (C). Graph depicts MFI of CD69 surface expression on B cells as determined via flow cytometry. **K-O.** Experiments were performed as in **H-J**, but *Egr2* transcript and EGR2 protein assayed instead via qPCR and intra-cellular staining respectively. **P.** Experiment performed as in (C). Graph depicts MFI of intracellular IRF4 protein expression in B cells as determined via flow cytometry. Data in this figure depict N=3 biological replicates for all panels except (D, K) as noted in Fig 7. Mean +/- SEM displayed for all graphs. Statistical significance was assessed with student's t-test with Holm-Sidak (A-C, D, E, J-L, O, P); two-way ANOVA with Tukey's (F-I, M, N). *p<0.05, **p<0.01, ***p<0.001, ****p<0.0001

**Supplemental figure 8. A.** Experiment performed as in Fig 7P. Graph depicts MFI of intracellular c-MYC expression on B cells as determined via flow cytometry.
**B.** Experiment performed as in Fig 7R. Graph depicts MFI of intracellular c-MYC expression on B cells as determined via flow cytometry. **C.** Experiment performed as in Fig S7C with competitive chimeras described in Fig S4M, N. Graph depicts MFI of intracellular c-MYC expression on B cells as determined via flow cytometry. **D.** Lymphocytes from *Batf*+/+, *Batf*+/-, and *Batf*-/- mice were stimulated with the given

doses of anti-IgM for 24 hours. Graph depicts intracellular BATF expression in B cells as determined via flow cytometry. **E, F.** Samples correspond to those described in Fig 8I, J, cultured in the presence of 20ng/ml BAFF. **E.** Shown is the ratio of each CD45.2+ B cell genotype relative to co-cultured CD45.1+ WT B cells, normalized to the unstimulated condition. **F.** Graph depicts division index for each CD45.2+ genotype. Data in this figure depict N=3 biological replicates. Mean +/- SEM displayed for all graphs. Statistical significance was assessed with two-way ANOVA with Tukey's (A, B, E, F); student's t-test with Holm-Sidak (C). *p<0.05, **p<0.01, ***p<0.001, ****p<0.0001

**Supplemental figure 9. A.** Experiment performed as in Fig 7P. Graph depicts MFI of surface CD86 expression on B cells as determined via flow cytometry.
**B.** Experiment performed as in Fig S7C with competitive chimeras described in Fig S4M, N. Graph depicts MFI of surface CD86 expression on B cells as determined via flow cytometry. **C, D.** Lymphocytes harvested from CD45.1+ *Nr4a1+/+* and CD45.2+ *Nr4a1-/-* mice were mixed 1:1 and co-cultured with the given doses of anti-IgM for 24 h (C) or 48 h (D). Graphs depict MFI of ICAM-1 surface expression on each genotype of B cells as determined via flow cytometry. **E.** Graph depicts adoptive transfer data from experiments described in 9A, B and presented in Fig 9J with addition of unimmunized controls and CD40L-/- hosts that did not receive adoptive transferred OTII cells (shaded gray points). Data in this figure depict N=3 biological replicates except (E) where individual biological samples are plotted. Mean +/- SEM displayed for all other graphs. Statistical significance was assessed with two-way ANOVA with Tukey's (A); student's t-test with Holm-Sidak (B-D). *p<0.05, **p<0.01, ***p<0.001, ****p<0.0001

**Supplementary Table 1. qPCR primers**

| Gene | Fwd primer | Rev primer |
|---|---|---|
| *Batf* | cacagaaagccgacacccctt | gctgtttgatctctttgcgga |
| *Ccl3* | accatgacactctgcaacca | cgatgaattggcgtggaatct |
| *Ccl4* | caaacctaaccccgagcaac | agaaacagcaggaagtggga |
| *Cd69* | agctcttcacatctggagagag | ccactattaacacagcccaagg |
| *Egr2* | ttgaccagatgaacggagtg | cagagatgggagcgaagcta |
| *Gfp* | cacatgaagcagcacgactt | tccttgaagtcgatgccctt |
| *cMyc* | tgaagttcacgttgagggg | agagctcctcgagctgtttg |
| *Nr4a1* | gcctagcactgccaaattg | ggaaccagagagcaagtcat |
| *Nr4a2* | tgaatgaagagagcggacaa | tgtcgtaattcagcgaagga |
| *Nr4a3* | aaacttgcagagcctgaacc | ctggtggtcctttaagctgc |

Supplemental Figure 1

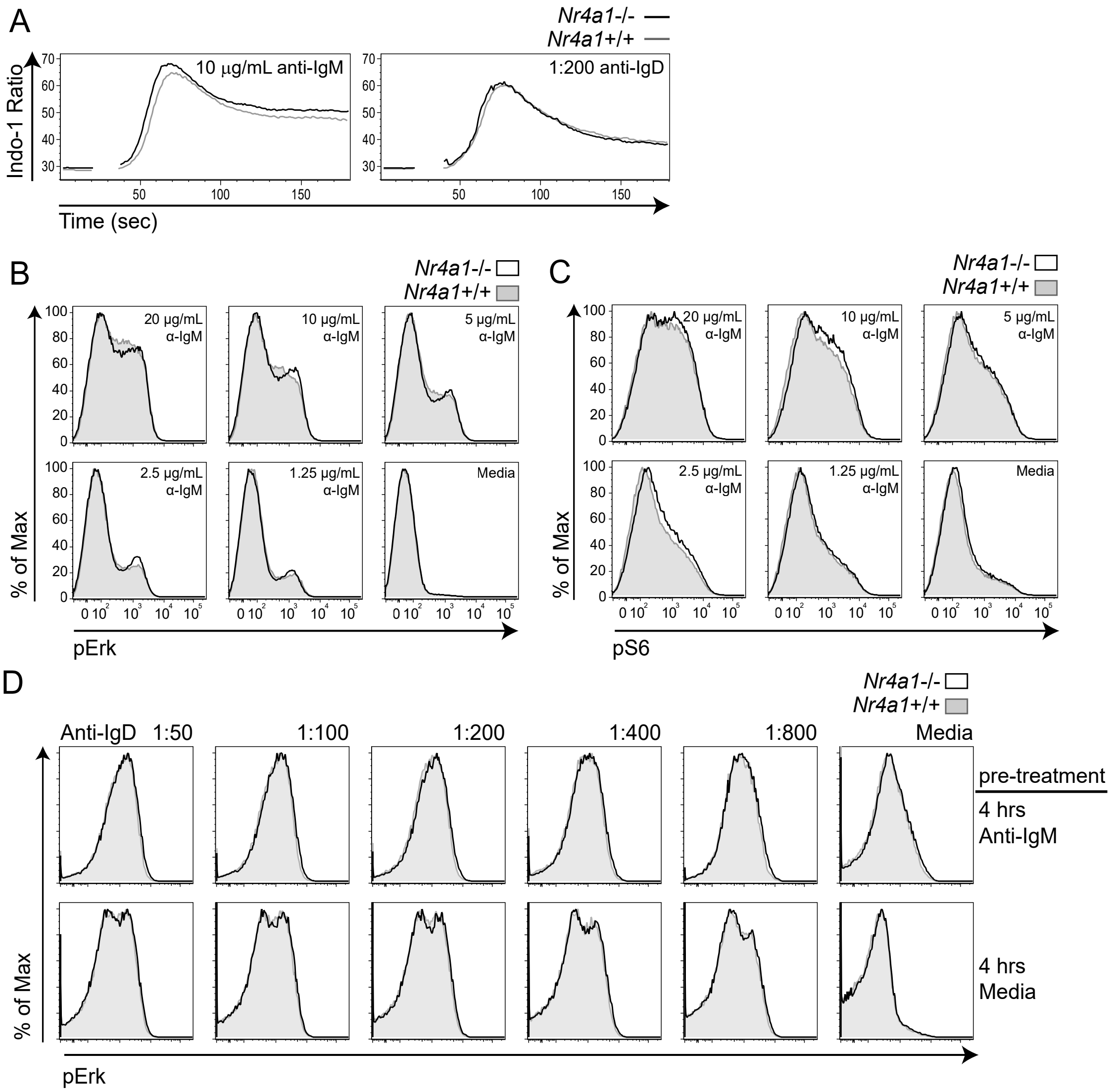

Supplemental Figure 2

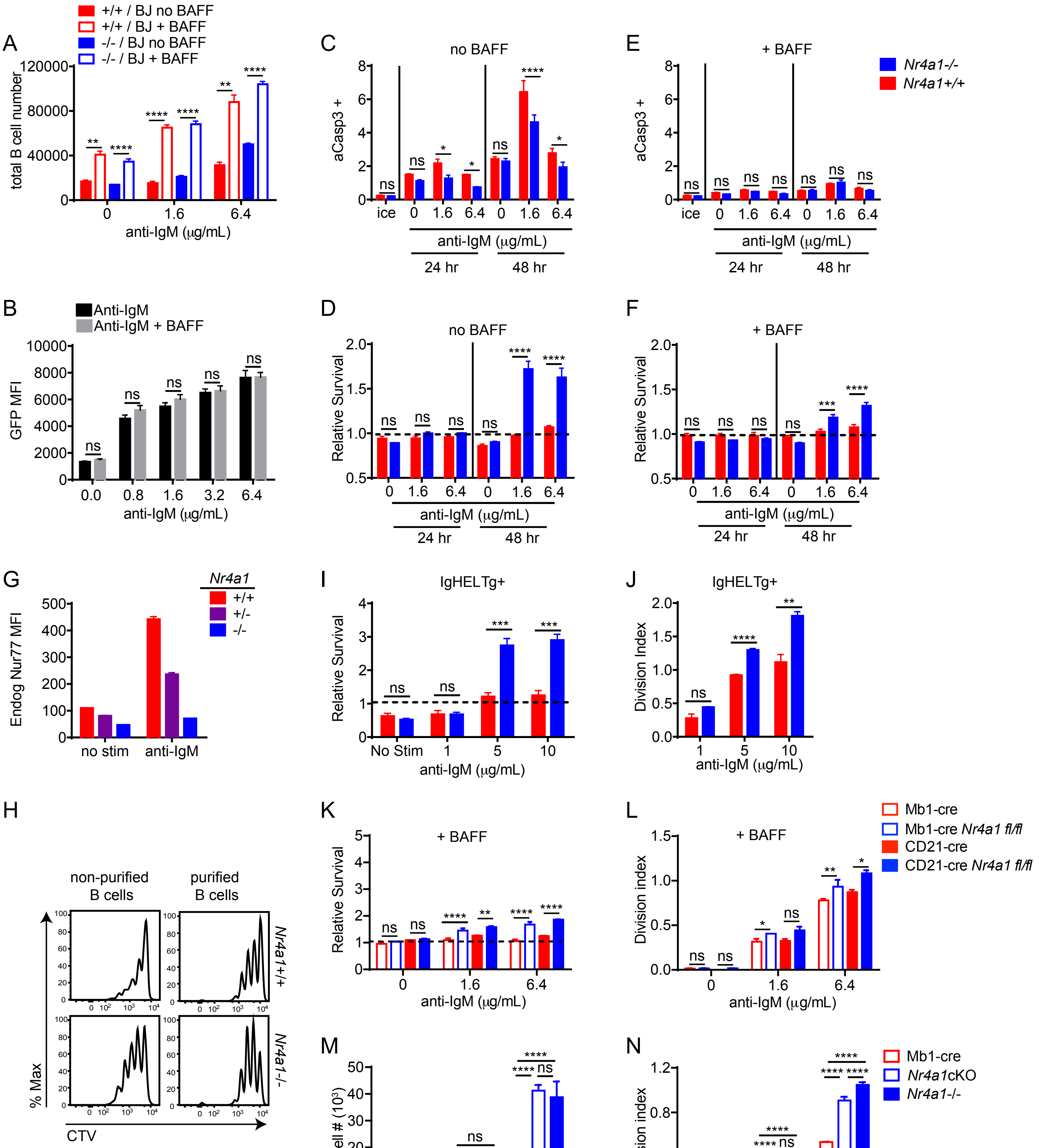

### Supplemental Figure 3

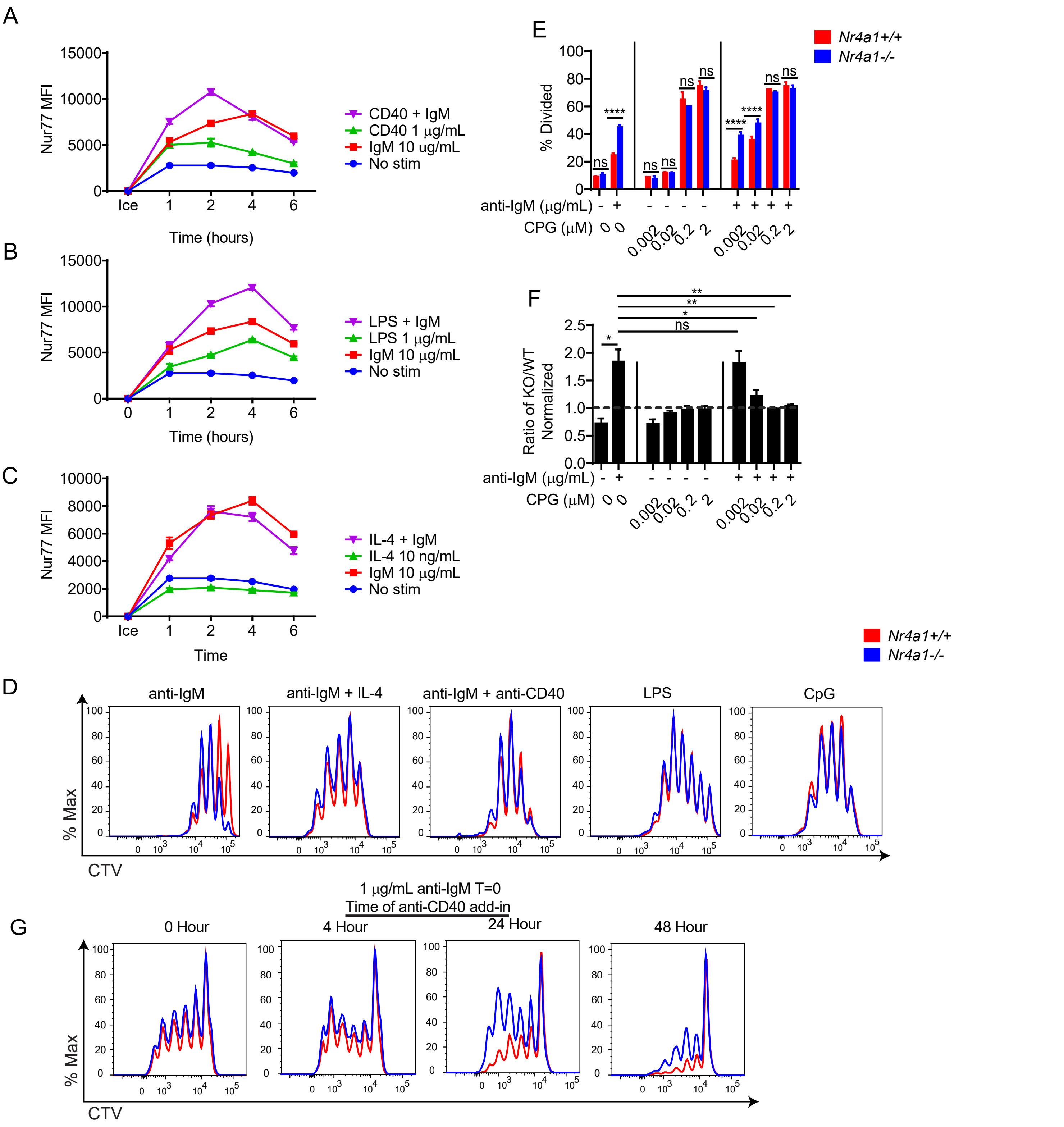

### Supplemental Figure 4

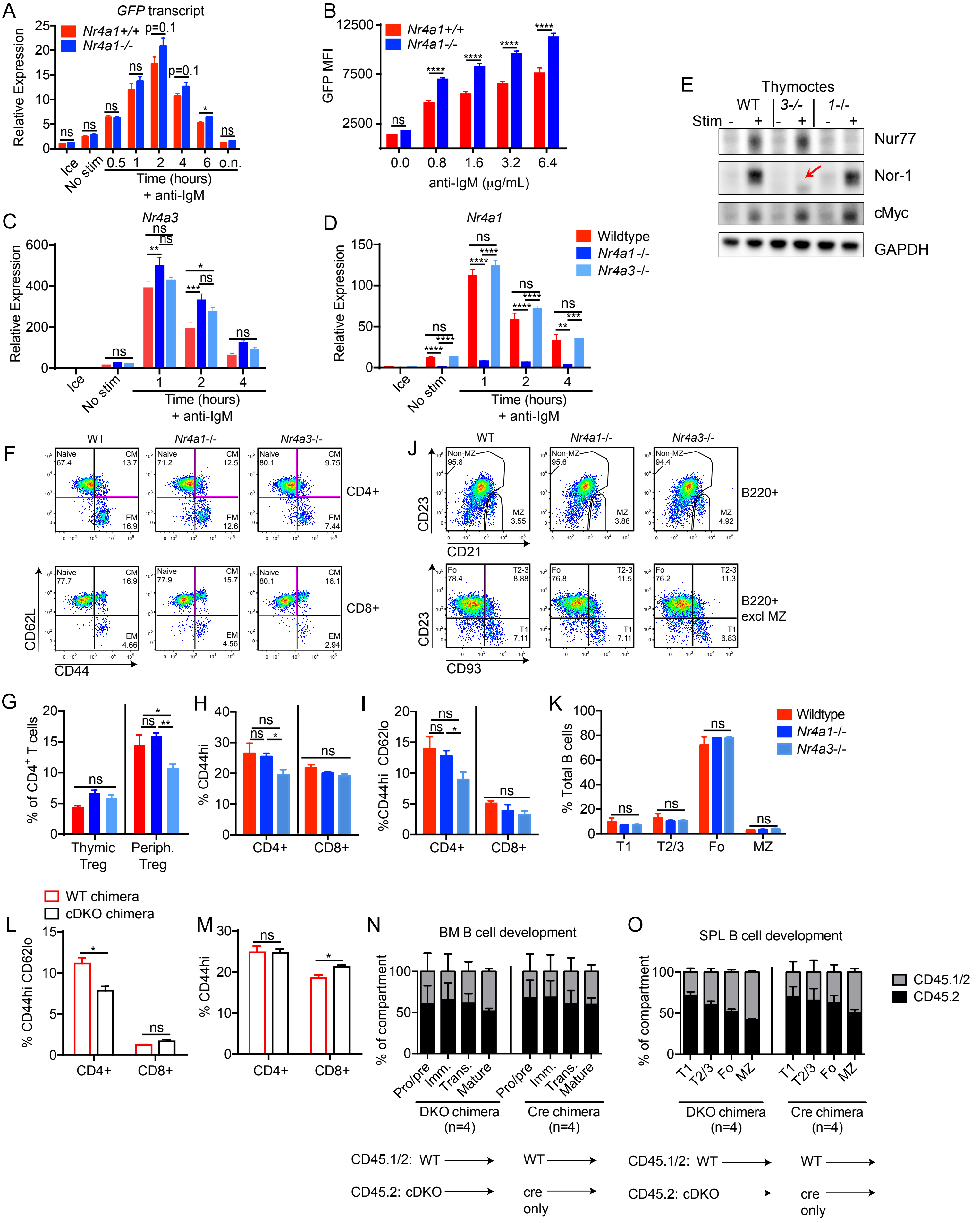

Supplemental Figure 5

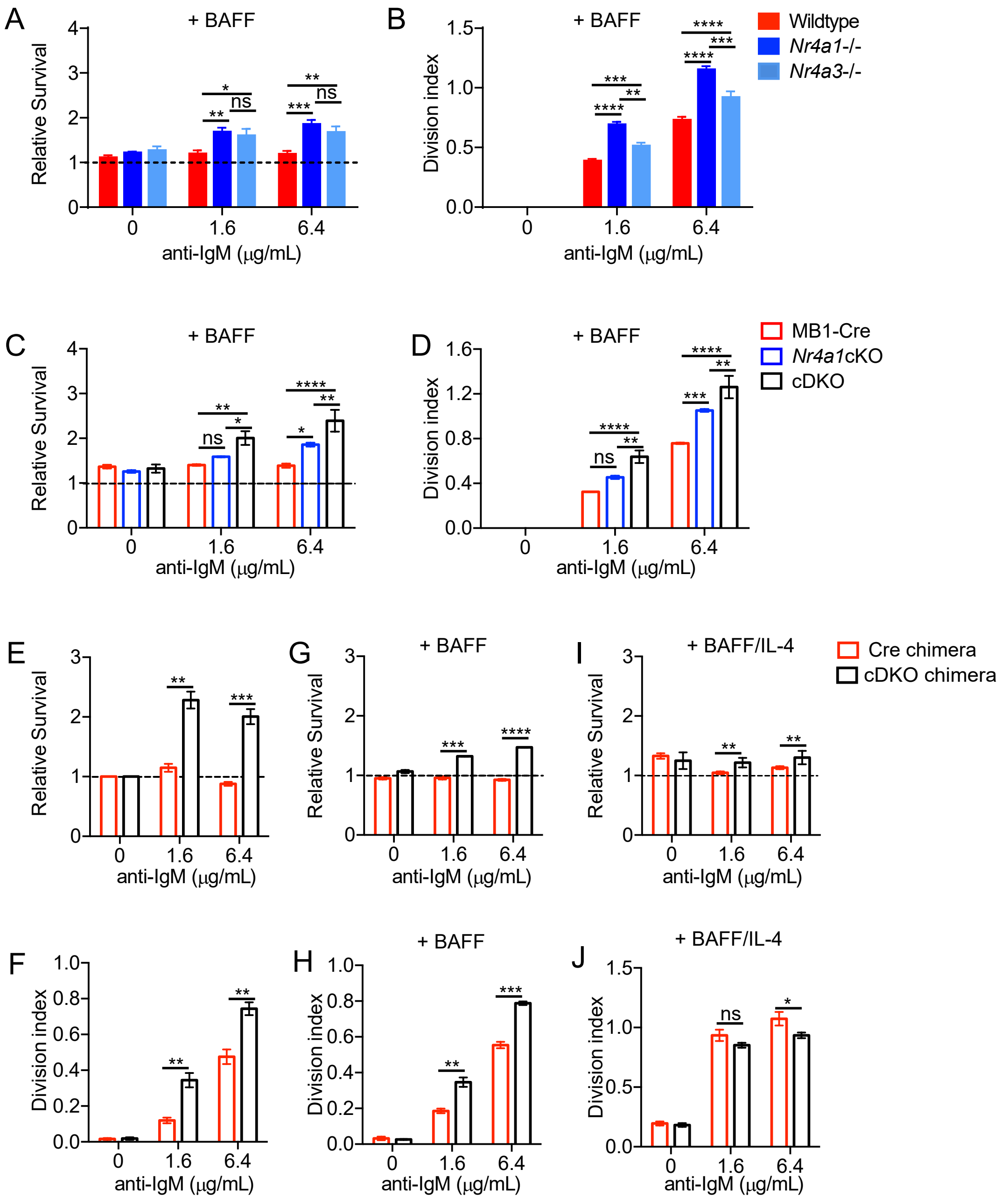

### Supplemental Figure 6

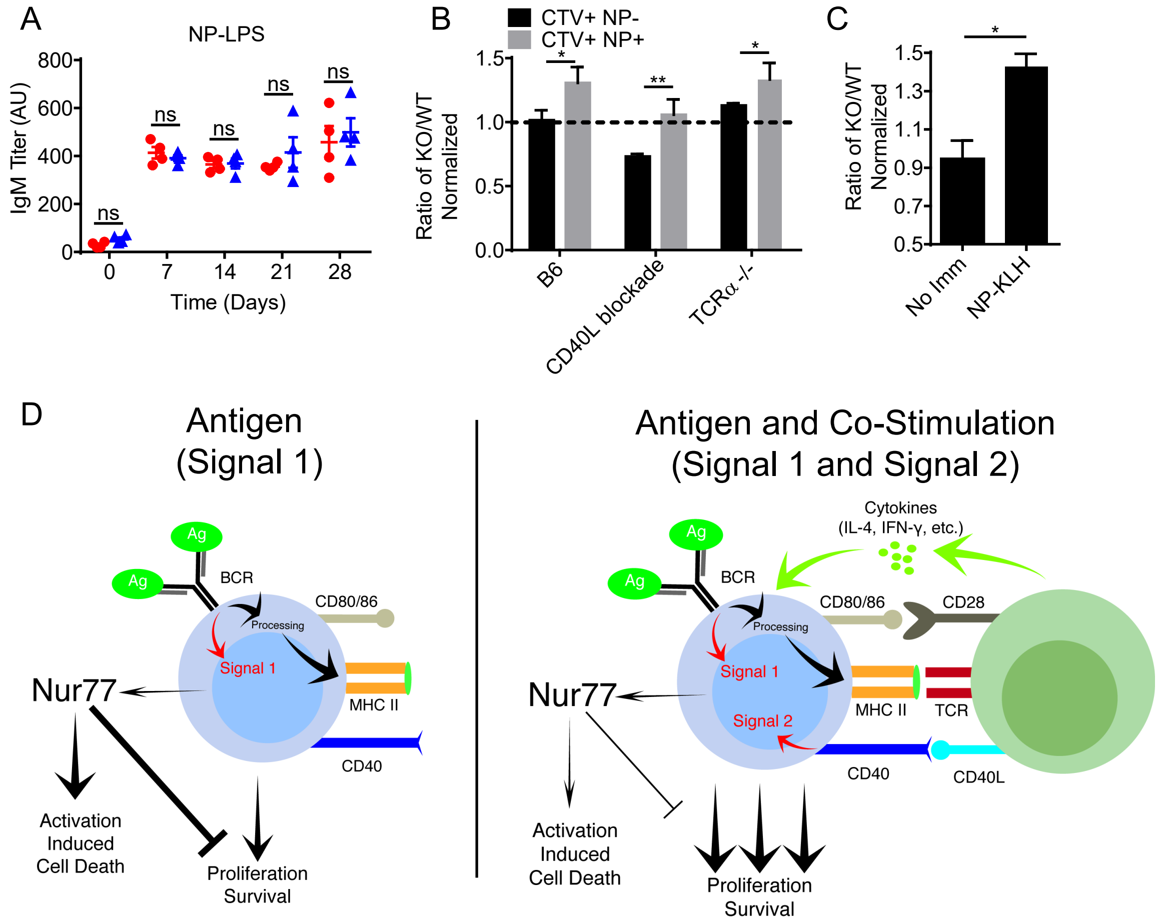

### Supplemental Figure 7

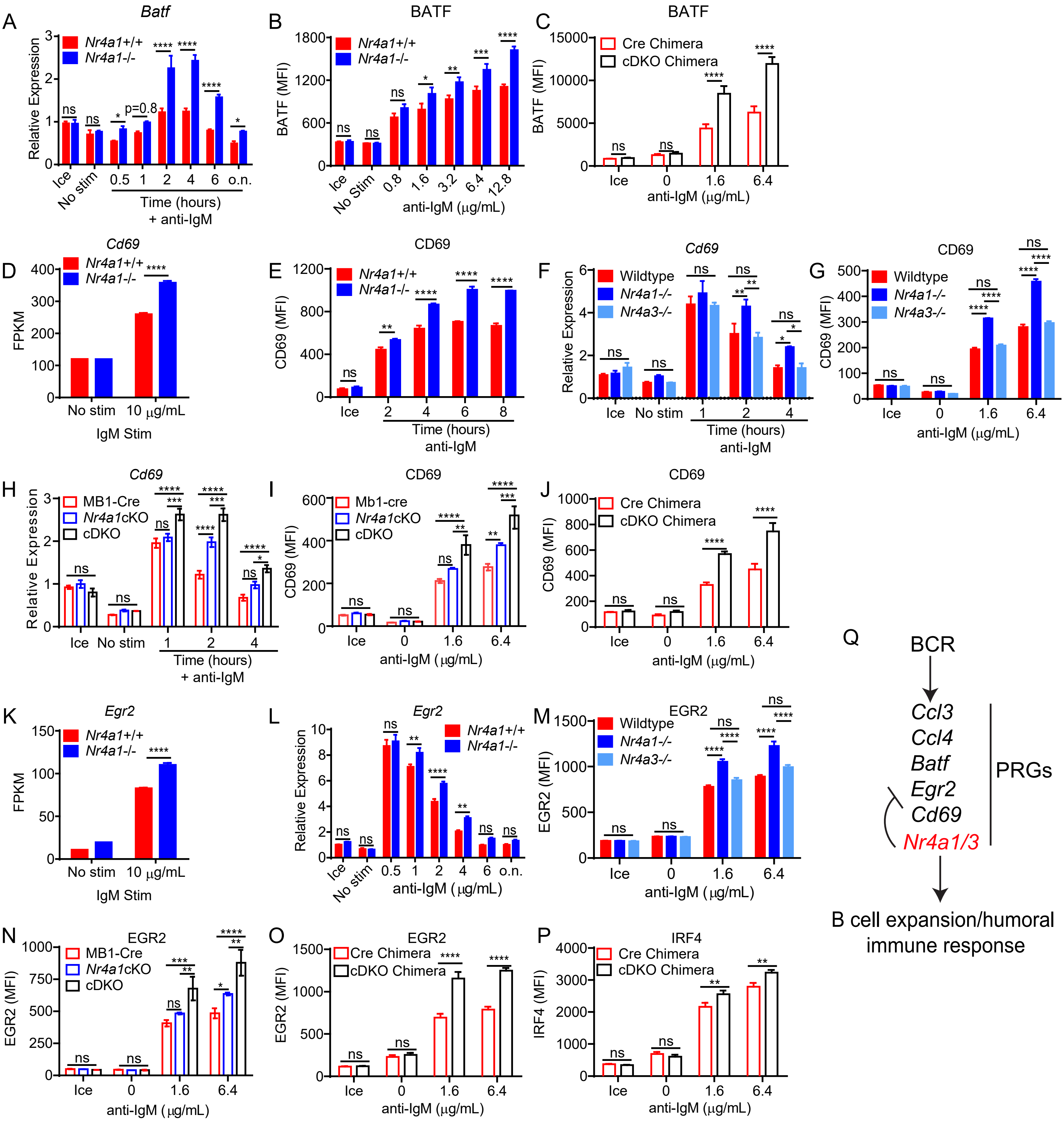

### Supplemental Figure 8

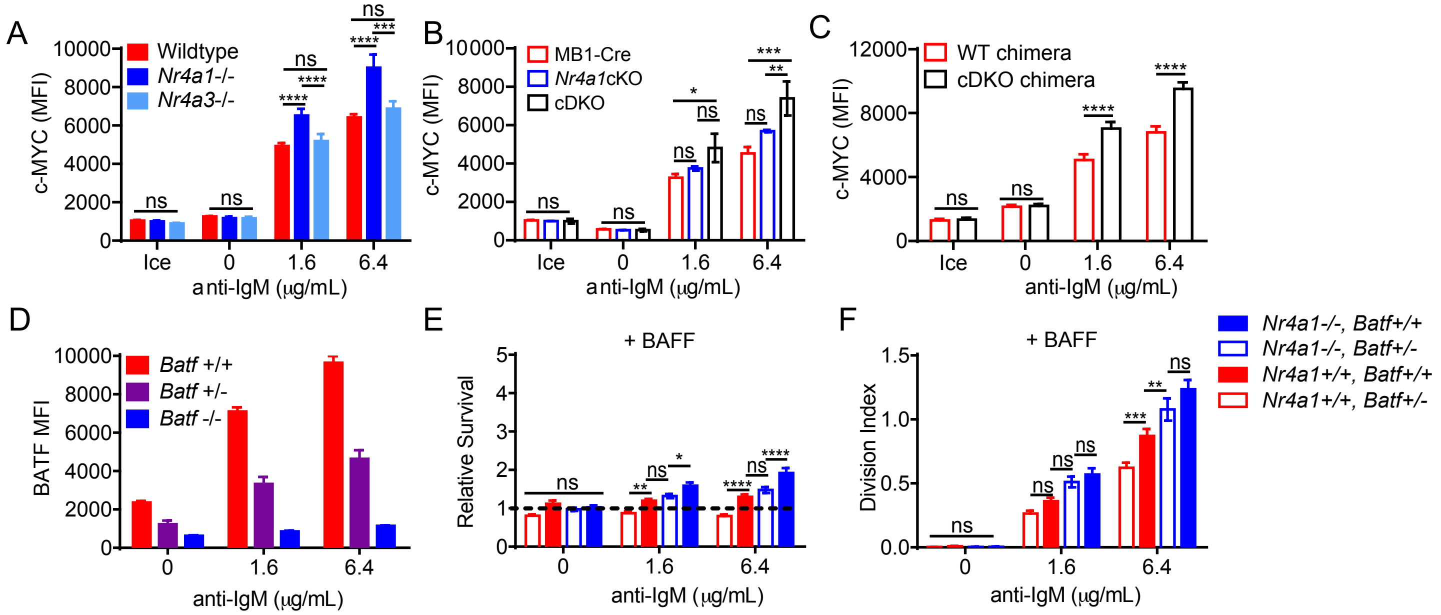

Supplemental Figure 9

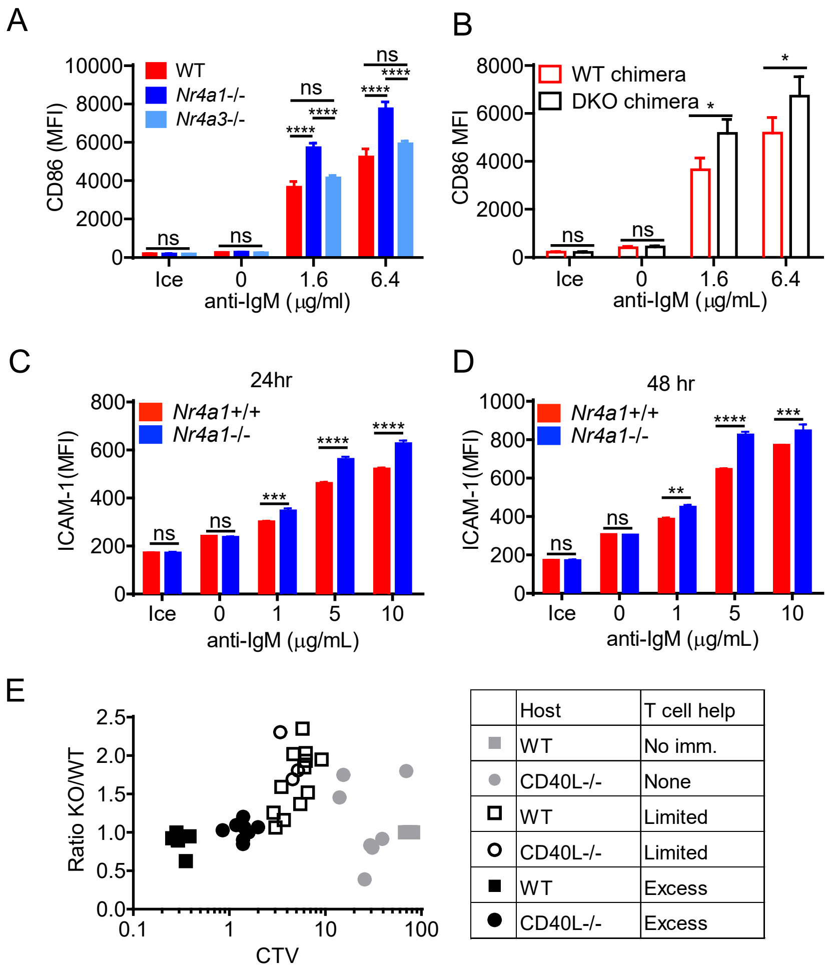


| | Host | T cell help |
|---|---|---|
| ■ (grey) | WT | No imm. |
| ● (grey) | CD40L-/- | None |
| □ | WT | Limited |
| ○ | CD40L-/- | Limited |
| ■ | WT | Excess |
| ● | CD40L-/- | Excess |
